# A high-affinity human PD-1/PD-L2 complex informs avenues for small-molecule immune checkpoint drug discovery

**DOI:** 10.1101/786319

**Authors:** Shaogeng Tang, Peter S. Kim

## Abstract

Immune checkpoint blockade of programmed death-1 (PD-1) by monoclonal antibody drugs has delivered breakthroughs in the treatment of cancer. Nonetheless, small-molecule PD-1 inhibitors could lead to increases in treatment efficacy, safety, and global access. While the ligand-binding surface of apo-PD-1 is relatively flat, it harbors a striking pocket in the murine PD-1/PD-L2 structure. An analogous pocket in human PD-1 may serve as a small-molecule drug target, but the structure of the human complex is unknown. Because the CC′ and FG loops in murine PD-1 adopt new conformations upon binding PD-L2, we hypothesized that mutations in these two loops could be coupled to pocket formation and alter PD-1’s affinity for PD-L2. Here, we conducted deep mutational scanning in these loops and used yeast surface display to select for enhanced PD-L2 binding. A PD-1 variant with three substitutions binds PD-L2 with an affinity two orders of magnitude higher than that of the wild-type protein, permitting crystallization of the complex. We determined the X-ray crystal structures of the human triple-mutant PD-1/PD-L2 complex and the apo triple-mutant PD-1 variant at 2.0 Å and 1.2 Å resolution, respectively. Binding of PD-L2 is accompanied by formation of a prominent pocket in human PD-1, as well as substantial conformational changes in the CC′ and FG loops. The structure of the apo triple-mutant PD-1 shows that the CC′ loop adopts the ligand-bound conformation, providing support for allostery between the loop and pocket. This human PD-1/PD-L2 structure provide critical insights for the design and discovery of small-molecule PD-1 inhibitors.

**Significance Statement:** Immune checkpoint blockade of programmed death-1 (PD-1) by monoclonal antibody drugs has transformed the treatment of cancer. Small-molecule PD-1 drugs have the potential to offer increased efficacy, safety, and global access. Despite substantial efforts such small-molecule drugs have been out of reach. We identify a prominent pocket on the ligand-binding surface of human PD-1 that appears to be an attractive small-molecule drug target. The pocket forms when PD-1 is bound to one of its ligands, PD-L2. Our high-resolution crystal structure of the human PD-1/PD-L2 complex facilitates virtual drug-screening efforts and opens additional avenues for the design and discovery of small-molecule PD-1 inhibitors. Our work provides a strategy that may enable discovery of small-molecule inhibitors of other “undruggable” protein-protein interactions.

## Introduction

Immune checkpoint blockade of programmed death 1 (PD-1) and its ligand 1 (PD-L1) has dramatically increased progression-free survival for many cancers (1-3). The first time that the FDA approved a cancer treatment based on a genetic biomarker rather than the primary site of origin was in 2017, when the monoclonal antibody (mAb) drug, pembrolizumab (Keytruda^®^), received approval for use in patients with microsatellite instability-high or mismatch repair deficient solid tumors (4, 5). Indeed, mAb drugs inhibiting immune checkpoints have ushered in an exciting new chapter in oncology.

Nevertheless, there is a desire for small-molecule inhibitors of immune checkpoints. First, in general, small molecules are expected to penetrate more effectively than mAbs into the tumor microenvironment, perhaps enhancing efficacy (6). In addition, if penetration into the brain is desired, small molecules can be effective (7, 8). Second, there are rare but severe immune-related side effects of checkpoint inhibition that require immediate drug discontinuation (9-12). Since mAbs have long half-lives in the body (typically weeks) (13), the treatment of such severe immune-related side effects is primarily supportive. Small-molecule checkpoint inhibitors could offer convenient dosing (e.g., once per day) while allowing for prompt drug removal if desired (14). Finally, small-molecule immune checkpoint inhibitors would facilitate cancer treatment in low- and middle-income countries by reducing production costs and eliminating the need for refrigeration during transportation and storage, in contrast to mAbs (15). Despite substantial efforts (16, 17), there are no well-characterized small-molecule ligands for PD-1.

The ligand-binding surface of human PD-1 is generally flat, lacking pockets considered suitable for binding small molecules (18). However, upon binding to PD-L1, a modest cavity forms on the ligand-binding surface of PD-1 (19). A similar cavity forms in murine PD-1 upon binding of PD-L1 (20). Importantly, when murine PD-1 binds a different ligand, PD-L2 (21), this cavity extends (Figures 1A-B) to a volume comparable to that occupied by established small-molecule inhibitors (22-25). Unfortunately, this murine structure is insufficient to provide a structural model for the analogous pocket in the human PD-1/PD-L2 complex, as the human and murine PD-1 proteins share sequence identities of only 63% (26).

**Figure 1.**
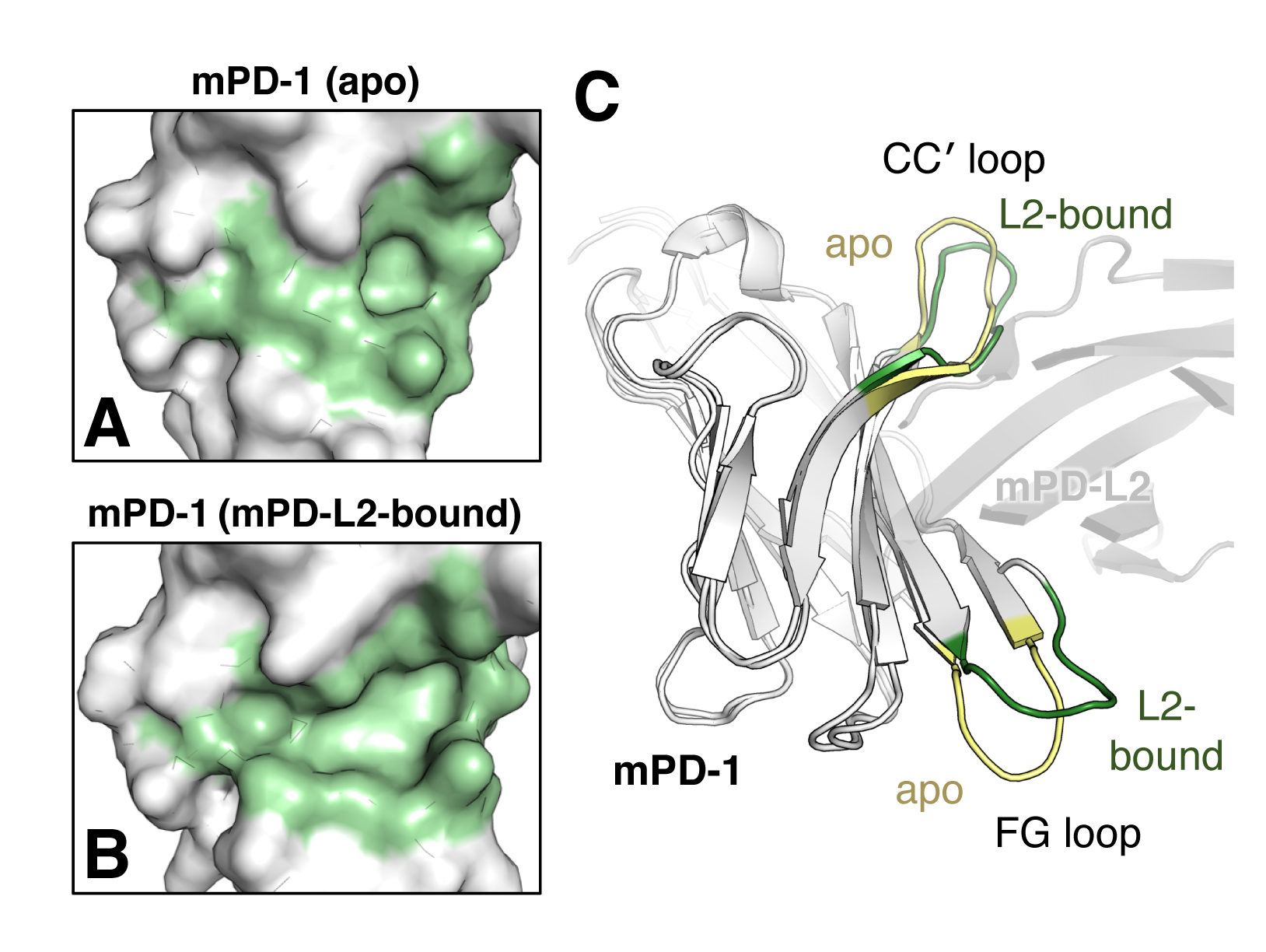
Conformational changes in murine PD-1 upon binding PD-L2. (A-B) Close-up views of space-filling models of (A) murine apo-PD-1 (PDB: 1NPU) and (B) murine PD-L2-bound murine PD-1 (PDB: 3BP5). The hydrophobic ligand-binding interface on PD-1 (pale green) forms a large pocket when murine PD-1 binds to PD-L2. (C) Overlay of ribbon diagrams of the murine apo-PD-1 (PDB: INPU) and the PD-L2-bound murine PD-1 (PDB: 3BP5). The CC loops and the FG loops adopt different conformations, and are highlighted for the apo (pale yellow) and PD-L2 bound (dark green) structures. mPD-1, murine PD-1; mPD-L2, murine PD-L2.

Although the murine PD-1/PD-L2 structure was determined over a decade ago (21), the structure of the human complex has not been reported. Our previous attempts to obtain diffraction-quality crystals of the human PD-1/PD-L2 complex were unsuccessful. Analyses of earlier structural studies (21, 26) revealed that formation of cavities on the ligand-binding surface of PD-1 is accompanied by changes in the structures of the CC′ and FG loops (Figure 1C). We therefore hypothesized that substitutions in these loops could have an allosteric effect on the conformations of PD-1 in the pocket region and alter its affinity for PD-L2. Using deep mutational scanning (27, 28) and yeast surface display (29), we selected for CC′ and FG loop variants of human PD-1 with enhanced PD-L2 binding. We identified a triple-mutant PD-1 that binds PD-L2 with nanomolar affinity and is amenable to crystallization, both alone and as a complex. The resulting X-ray crystal structures confirm that a prominent pocket forms in human PD-1 upon binding of PD-L2 and support the notion of allostery between the pocket and the CC′ and FG loops. The pocket identified here in human PD-1 can serve as a template for virtual drug discovery (30) and opens up additional avenues for the discovery of small-molecule PD-1 inhibitors.

## Results

### Engineering human PD-1 loop variants with enhanced PD-L2 affinity

Substantial efforts by us and others (31) to crystallize the human PD-1/PD-L2 complex were previously unsuccessful. Earlier studies (18, 19, 21) indicated that the PD-1 ligand-binding interface consists of a hydrophobic core, the CC′ loop, and the FG loop (Figure 2A), and that formation of a complex with ligands results in loop movement and pocket formation in the hydrophobic core. We hypothesized that mutations in these two loops of PD-1 were coupled to pocket formation and could alter PD-1’s affinity for PD-L2. Consistent with this hypothesis, we found that poly-glycine mutants of these loops in human PD-1 substantially decreased affinities for PD-L2 (Figures S1A-B).

**Figure 2.**
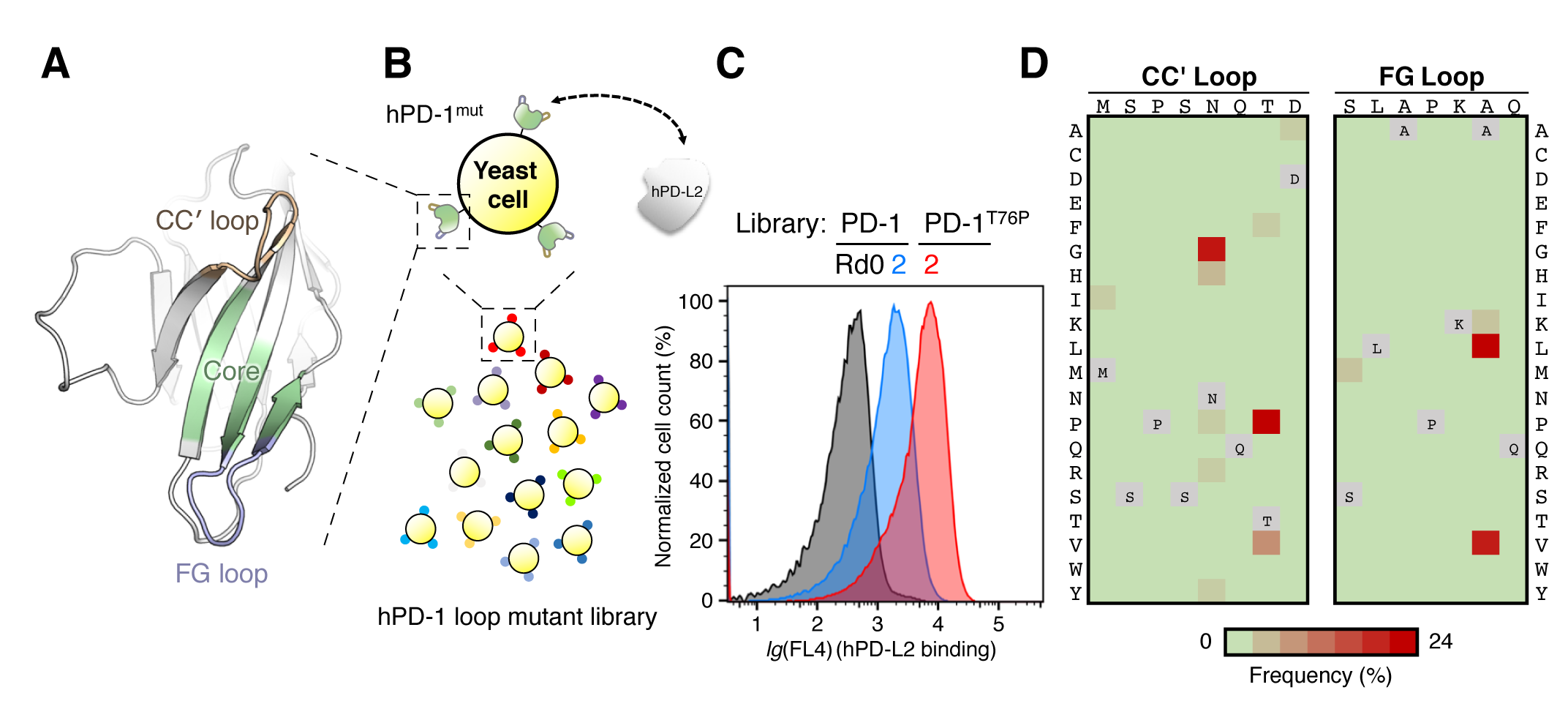
Engineering PD-1 loop variants with enhanced PD-L2 affinity. (A) Ribbon diagram of the human PD-1 ectodomain, highlighting the CC loop (wheat), the FG loop (light blue), and the hydrophobic ligand-binding interface (pale green). (B) Schematic of yeast surface display of a human PD-1 (hPD-1) loop variant library (colored spheres) and selection for binding of a recombinant human PD-L2 (hPD-L2) ectodomain. (C) Overlay of flow-cytometric histograms of the PD-1 loop-variant yeast library at selection rounds 0 (black) and 2 (blue), and the PD-1^T76P^ loop variant yeast library at selection round 2 (red). Yeast cells were stained with 10 nM PD-L2-Fc, followed by Alexa Fluor 647-labeled secondary antibody against human Fc. Yeast cells exhibit enhanced PD-L2-Fc binding after rounds of selection. (D) Frequency heatmaps of human PD-1 amino acid substitutions in the CC loop (left) and the FG loop (right) after selection round 2 of the PD-1 loop-variant yeast library using PD-L2-Fc. Substitutions of N74G and T76P were identified in the CC loop, and A132V and A132L in the FG loop.

Since we were particularly interested in the structure of the PD-1 pocket when bound to PD-L2, we maintained residues in the hydrophobic core and performed directed evolution exclusively in the CC′ loop (residues 70-78) and the FG loop (residues 127-133) of human PD-1. We used deep mutational scanning (27, 28) to construct loop-variant libraries with trinucleotides encoding each of 20 residues at each position. We next used yeast surface display (29) (Table S1) with a recombinant human PD-L2-human Fc fusion protein as the selection agent (Figure 2B). After two rounds of selection using magnetic- and fluorescence-activated cell sorting (MACS and FACS) (Figure 2C), we isolated human PD-1 loop-variant clones with single-residue substitutions. Substitutions at two residues were identified in the CC′ loop (N74G and T76P) and at one residue in the FG loop (A132V, A132L) (Figure 2D). In contrast, when we used the same yeast library and selected with PD-L1-Fc, we only isolated the A132 substitutions as high-affinity variants (Figures S1C-E), suggesting that the N74G and T76P variants are PD-L2-binding specific. We chose PD-1^T76P^ as a template to generate a second PD-1 loop variant library and selected for further enhancement of PD-L2 binding (Figure 2C). As a result, we obtained a PD-1 triple mutant (Figures S1F-G) that contains all three substitutions identified from the first library: N74G, T76P, and A132V.

### PD-1 loop variants showed increased binding affinity and association kinetics for PD-L2 and PD-L1

To validate the detected enhancement in affinity, we recombinantly expressed and purified human PD-1 and the loop variants, as well as the human PD-L2 and PD-L1 ectodomain proteins. Using bio-layer interferometry, we compared the binding of PD-L2 to wild-type PD-1 and to the variants (Figures 3A and S2A). Wild-type PD-1 binds PD- L2 with a *KD* of 500 nM; the variants all exhibit increased PD-L2 affinity, with *KD* values of 170 nM for N74G, 12 nM for T76P, and 69 nM for A132V (Figure 3C). Remarkably, the PD-1 triple mutant has a *KD* of 2.6 nM for PD-L2, constituting a ∼200-fold increase in affinity (Figure 3C). The triple mutant also shows substantially higher affinity for PD-L1 (Figures 3B-C). The A132V mutant has higher affinity for PD-L1, consistent with previous reports (21, 31-33), but the N74G and T76P single mutants have minor effects (Figures 3B-C and S2B-C). Thus, this human PD-1 triple mutant exhibits a potent enhancement of binding affinity for both PD-L1 and PD-L2.

**Figure 3.**
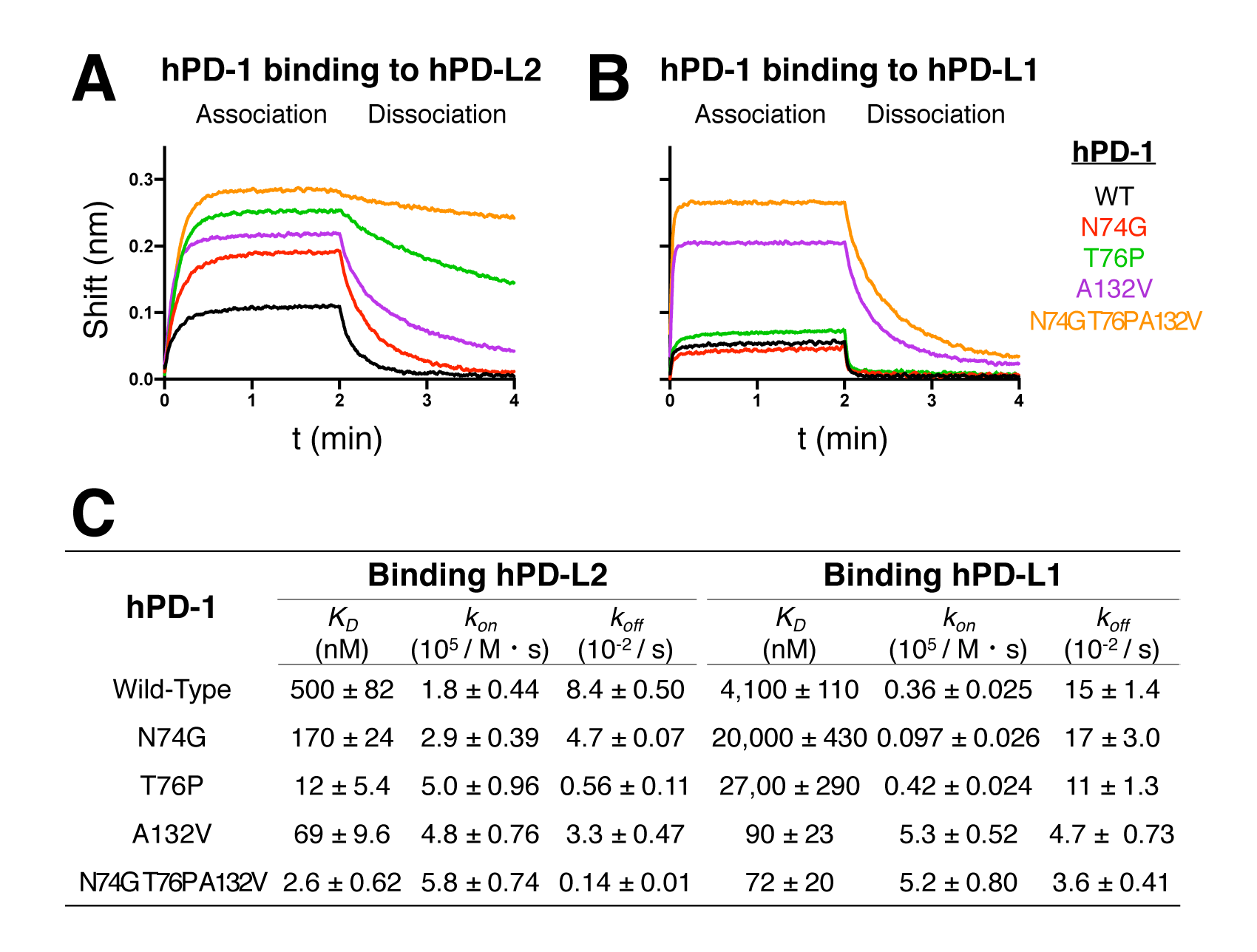
PD-1 loop variants with increased affinity and association kinetics for PD-L2 and PD-L1. (A-B) Binding of sensor-loaded PD-1 and the loop variants to (A) 190 nM PD-L2 and (B) 1.1 μM PD-L1 using bio-layer interferometry. Corresponding PD-1-Fc proteins were loaded on anti-human IgG Fc capture (AHC) biosensors. Association and dissociation were each monitored for 2 min. (C) Summary of binding affinity (*K_D_*) and kinetic parameters (association constant *k_on_,* dissociation constant *k_off_*) for the PD-1 loop variants binding to PD-L2 or PD-L1. Fitting of binding curves was performed in Graphpad Prism 8 using built-in equations of “Receptor binding - kinetics” model. Means and standard deviations were calculated from 3-4 independent experiments.

Bio-layer interferometry of ligand binding also enabled us to determine association constants (*kon*). Compared to wild-type PD-1, all loop variants showed increased *kon* for binding PD-L2 (Figure 3C). The PD-1 triple mutant underwent a 3-fold increase of *kon* for PD-L2, and a 14-fold increase for PD-L1 (Figure 3C). These results suggest that these amino acid substitutions in the loops stabilize the ligand-bound state among the conformational ensembles of apo-PD-1 ((34); see however, (35)).

### X-ray crystal structure of the human PD-1/PD-L2 complex

We then attempted to crystalize the human PD-1/PD-L2 complex using the PD-1 triple mutant. Site-directed mutagenesis was used to remove all N-linked glycosylation sites in each protein in an effort to aid crystallization (Table S2). Co-expression of the PD-1 triple mutant and the immunoglobulin variable (IgV) domain of PD-L2 yielded a stable and 1:1 stoichiometric complex (Figures S3A-B). We successfully obtained crystals of the human PD-1^N74G T76P A132V^/PD-L2^IgV^ complex and determined an X-ray co-crystal structure at 2.0 Å resolution (Figures 4A, S3C-D). The crystal contains one PD-1/PD-L2 complex per asymmetric unit, with space group *P* 2_1_ 2_1_ 2_1_ (Table 1). This structure reveals that the human PD-1/PD-L2 complex adopts an overall architecture similar to that previously determined for the murine PD-1/PD-L2 complex (21) with a Cα root-mean-square deviation (R.M.S.D.) of 3.8 Å. To our knowledge, this human PD-1/PD-L2 structure is the first reported human PD-L2 structure.

**Figure 4.**
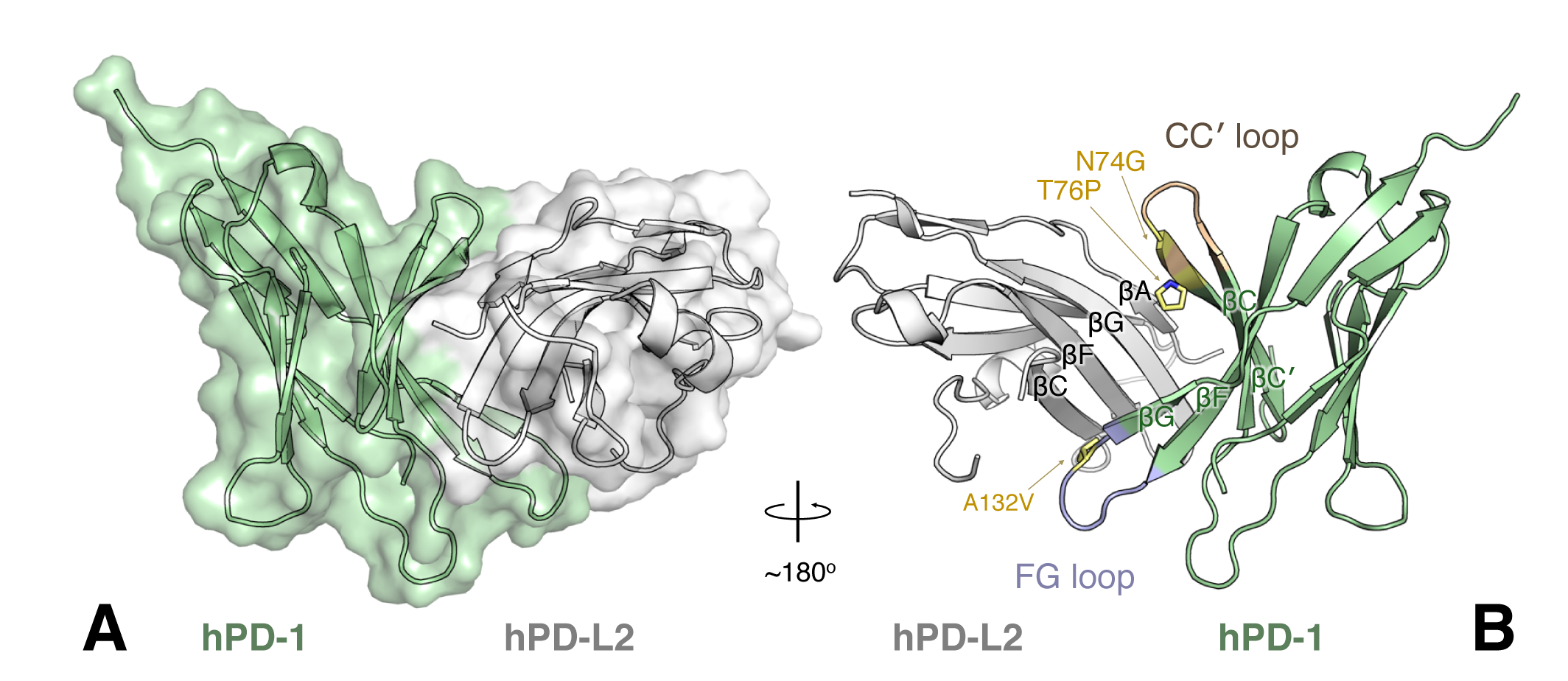
X-ray crystal structure of the human PD-1/PD-L2 complex. (A) Overlay of a space-filling diagram and a ribbon diagram of the complex of human PD-1^N74G T76P A132V^ (pale green) and PD-L2^IgV^ (grey), showing the overall architecture of the human PD- 1/PD-L2 complex. (B) Ribbon diagram of a ∼180° rotation view of (A) with the CC loop colored in wheat and the FG loop in light blue. The location of the substitutions of N74G, T76P, and A132V are labeled and their sidechains are indicated with sticks (pale yellow). The β-sheets on the interacting faces of each protein are labeled.

**Table 1.** Crystallographic data collection and refinement statistics

|  | **PD1<sup>N74G T76P A132V</sup>**<br>**/ PD-L2<sup>IgV</sup>** | **Apo-**<br>**PD1<sup>N74G T76P A132V</sup>** | **Apo-**<br>**PD1<sup>T76P A132V</sup>** |
| --- | --- | --- | --- |
| Wavelength (Å) | 0.978 | 0.978 | 0.978 |
| Resolution range (Å) | 37.5 - 1.99<br>(2.06 - 1.99) | 36.5 - 1.18<br>(1.23 - 1.18) | 36.5 - 1.42<br>(1.48 - 1.42) |
| Space group | *P* 2<sub>1</sub> 2<sub>1</sub> 2<sub>1</sub> | *P* 3<sub>2</sub> 2 1 | *P* 3<sub>2</sub> 2 1 |
| Unit cell | 41.3 67.8 89.7<br>90 90 90 | 46.2 46.2 89.3<br>90 90 120 | 46.2 46.2 89.4<br>90 90 120 |
| Total reflections | 185797 (11081) | 400313 (24984) | 171335 (11683) |
| Unique reflections | 17750 (1645) | 36661 (3544) | 21301 (2090) |
| Multiplicity | 10.4 (6.7) | 10.9 (7.0) | 8.0 (5.6) |
| Completeness (%) | 98.6 (90.6) | 99.7 (98.8) | 99.7 (98.2) |
| Mean I/sigma(I) | 16.1 (2.28) | 28.5 (2.79) | 23.3 (2.40) |
| Wilson B-factor | 35.8 | 16.7 | 21.9 |
| *R*<sub>merge</sub> | 0.139 (0.723) | 0.0521 (0.539) | 0.0903 (1.03) |
| *CC*<sub>1/2</sub> | 0.992 (0.780) | 0.999 (0.856) | 0.998 (0.769) |
| *CC*<sup>*</sup> | 0.998 (0.936) | 1.00 (0.960) | 0.999 (0.932) |
| *R*<sub>work</sub> | 0.196 (0.292) | 0.154 (0.192) | 0.158 (0.193) |
| *R*<sub>free</sub> | 0.226 (0.339) | 0.164 (0.233) | 0.189 (0.263) |
| Number of non-hydrogen atoms | 1782 | 1156 | 1143 |
| macromolecules | 1654 | 1001 | 1056 |
| water | 127 | 144 | 82 |
| Protein residues | 210 | 112 | 116 |
| RMS (bonds) (Å) | 0.013 | 0.009 | 0.016 |
| RMS (angles) (°) | 1.48 | 1.35 | 1.60 |
| Ramachandran favored (%) | 99 | 100 | 99 |
| Ramachandran outliers (%) | 0 | 0 | 0 |
| Clashscore | 8.32 | 0.99 | 5.66 |
| Average B-factor | 50.8 | 23.4 | 30.3 |
| macromolecules | 50.6 | 21.1 | 30.9 |
| solvent | 53.8 | 38.2 | 39.1 |

Statistics for the highest-resolution shell are shown in parentheses.

The human PD-1/PD-L2 interface is formed by the front β-sheets of both IgV domains (Figure 4B), burying 1,840 Å^2^ (14% of the total) of the solvent-accessible surface area. In the interface, notable interacting residues include the three highly conserved aromatics W110_L2_, Y112_L2_, and Y114_L2_ from βG of the PD-L2 IgV domain. The sidechains of these residues point into the center of the PD-1 ligand-binding surface (Figures S4A-B). To validate the interactions observed at the PD-1/PD-L2 interface, we performed site-directed mutagenesis on several PD-1 and PD-L2 interfacial residues. Bio-layer interferometry revealed reduced binding of PD-1 interface mutants to PD-L2, and PD-L2 interface mutants to PD-1 (Figures S4C-D), consistent with our co-crystal structure. The high-affinity loop substitutions of PD-1 localize to the interface (Figure 4B). Among them, T76P and A132V make additional contacts to PD-L2 that likely contribute to the increase in affinity (Figures S4E-H).

### X-ray crystal structures of human apo-PD-1 loop variants

To assist analyses of the conformational changes in PD-1 associated with PD-L2 binding, we crystallized two human apo-PD-1 loop variants (Table S2) and determined X-ray crystal structures at 1.2 Å and 1.4 Å resolution for PD-1^N74G T76P A132V^ (Figures S5A, S5C) and PD-1^T76P A132V^ (Figures S5B, S5D), respectively. Crystals of both variants contain a single PD-1 molecule per asymmetric unit, with space group *P* 3_2_ 2 1 (Table 1). Both PD-1 variants were well defined by the electron density maps, with the notable exception of the CC′ loop discussed further below (Figures S5E-F). Superimposing the apo and PD-L2-bound PD-1^N74G T76P A132V^ structures resulted in a C_α_ R.M.S.D. of 1.6 Å.

The C′D loop of PD-1 (residues 83-92) is a major part of the pembrolizumab epitope (36-38). This loop is not resolved in earlier structures of human PD-1 in the absence of pembrolizumab (19, 31, 39), but is clear in both of our apo-PD-1 structures. Our results indicate that the conformation of the loop changes substantially upon antibody binding (Figure S5G).

### Formation of a prominent pocket in human PD-1 upon binding PD-L2 with an architecture distinct from the murine pocket

Our crystal structures of the human PD-1/PD-L2 complex and apo-PD-1 variants allowed us to examine formation of the human PD-1 pocket in the PD-1/PD-L2 interface. Although the human apo-PD-1 variant has a flat ligand-binding interface (Figure 5A), there are rearrangements in this interface upon binding PD-L2. These rearrangements involve residues in βC (F63, V64, N66, Y68), βF (L122, G124, I126), βG (I134, E136), and the C′D loop (E84) to form a deep and extended pocket (Figure 5B). Each of these residues in PD-1 is within 4.4 Å of a PD-L2 residue (Figure S4I). This pocket accommodates PD-L2 sidechains including the aromatic residues W110L2 and Y112L2 (Figure 5C).

**Figure 5.**
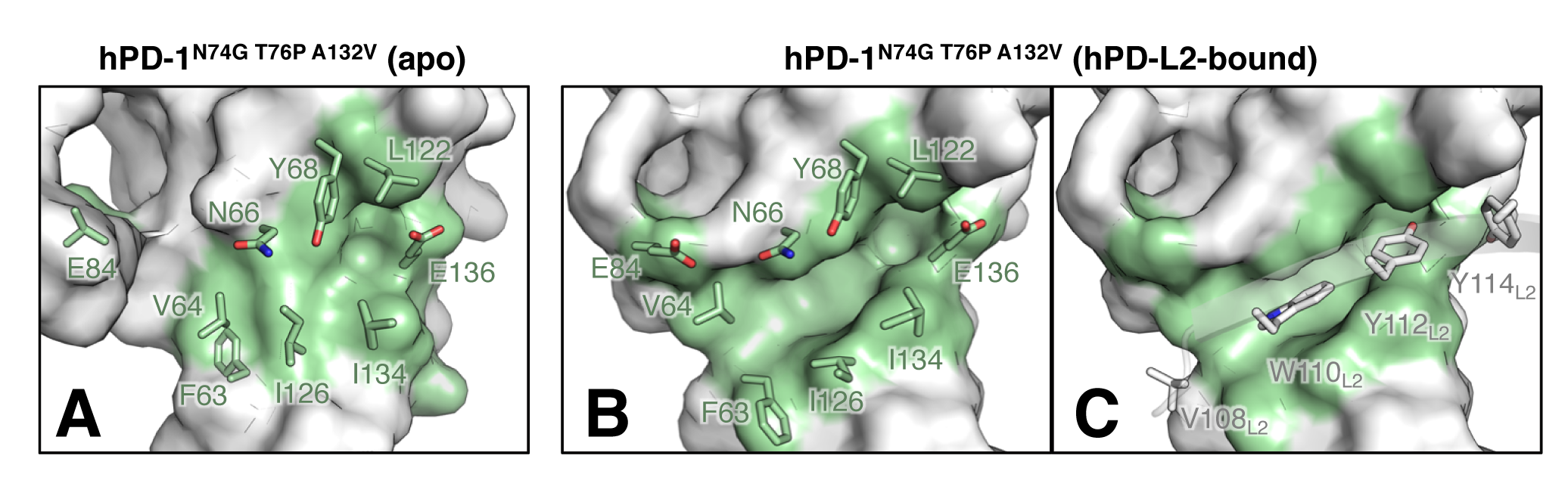
Formation of a prominent pocket in human PD-1 upon binding PD-L2. Close-up views of space-filling models of (A) apo-human PD-1^N74G T76P A132V^ and (B-C) human PD- L2-bound human PD-1^N74G T76P A132V^ overlaid with pocket residues shown as sticks. In (C) a ribbon diagram of the βG of PD-L2 is shown with PD-L2-interacting residues overlaid as sticks and labeled with an L2 subscript. A 170 Å^3^ funnel-shaped pocket forms (left, entrance; right, exit) when human PD-1 binds PD-L2.

Comparison of the PD-1 pockets in the human and murine PD-1/PD-L2 complexes revealed striking differences in pocket geometries. The human pocket adopts an open, funnel-shaped architecture. Compared to the murine pocket (Figures 1B, S6B-C), the human pocket has a wider entrance and a narrower exit (Figure 5B). The distinct pocket geometries arise from at least two factors. First, human PD-1 employs a different subset of interfacial residues to form the pocket than the murine version. Human PD-1 lacks an ordered βC′′ strand and thus the open pocket is formed by rearranging residues F63, V64, and E84. In contrast, the murine pocket is closed, with sidechains of A81 and S83 forming a boundary (Figures S6A-B). Second, several sequence variations exist among the residues that form the pocket. For example, V64 and Y68 in human PD-1 are substituted with M64 and N68 in murine PD-1, respectively (Figures 5B and S6B). To quantitatively evaluate the pocket dimensions, we measured pocket volumes using POCASA 1.1 (40). The human and murine pockets have volumes of 170 Å^3^ and 220 Å^3^, respectively. Notably, these pockets are comparable in size to other protein cavities with established small-molecule inhibitors (160-800 Å^3^) (22-25).

We compared our human PD-1/PD-L2 structure (Figure S6E) with the previously determined human PD-1/PD-L1 structure (19) (Figure S6F). Superimposing the two structures resulted in a Cα R.M.S.D. of 1.5 Å for PD-1 residues. Binding PD-L1 triggers formation of a much smaller cavity in human PD-1, with a volume of 80 Å^3^ (Figure S6D). PD-L1 lacks a large aromatic sidechain corresponding to W110L2, so the PD-1 rearrangement only involves accommodation of a small subset of the interfacial residues, including the sidechain of Y123_L1_, which corresponds to PD-L2 residue Y112_L2_ (Figures S6E-H). These results indicate that the core of the human PD-1 interface has remarkable structural plasticity, with the ability to form pockets with varied dimensions to interact with different PD-1 ligands.

### The CC′ loop in triple-mutant PD-1 adopts a ligand-bound conformation in the absence of ligand

We also detected conformational changes in the CC′ and FG loops when human PD-1 binds PD-L2 (Figures 6A-B). Earlier studies reported that the CC′ loop undergoes a substantial conformational change when human PD-1 binds PD-L1 (19, 39). This CC′ loop conformational change is even larger in the human PD-1/PD-L2 structure reported here (Figures 6A and S7A). Strikingly, in the absence of ligands, the CC′ loop conformations of the PD-1 triple and double mutants resemble those of the ligand-bound conformations (Figure S7A). For example, a 4.8 Å shift occurs between the C_α_ of T76 and P76 in the PD-1 triple mutant of apo-PD-1 (Figure 6A). When the PD-1 triple mutant binds PD-L2, the sidechain of P76 maintains the same conformation as the unbound protein (Figure 6A). An increased population of the ligand-bound conformations of the mutant apo-PD-1 proteins is consistent with increased association constants (*kon*) of the PD-1 variants (Figure 3C and S2C).

**Figure 6.**
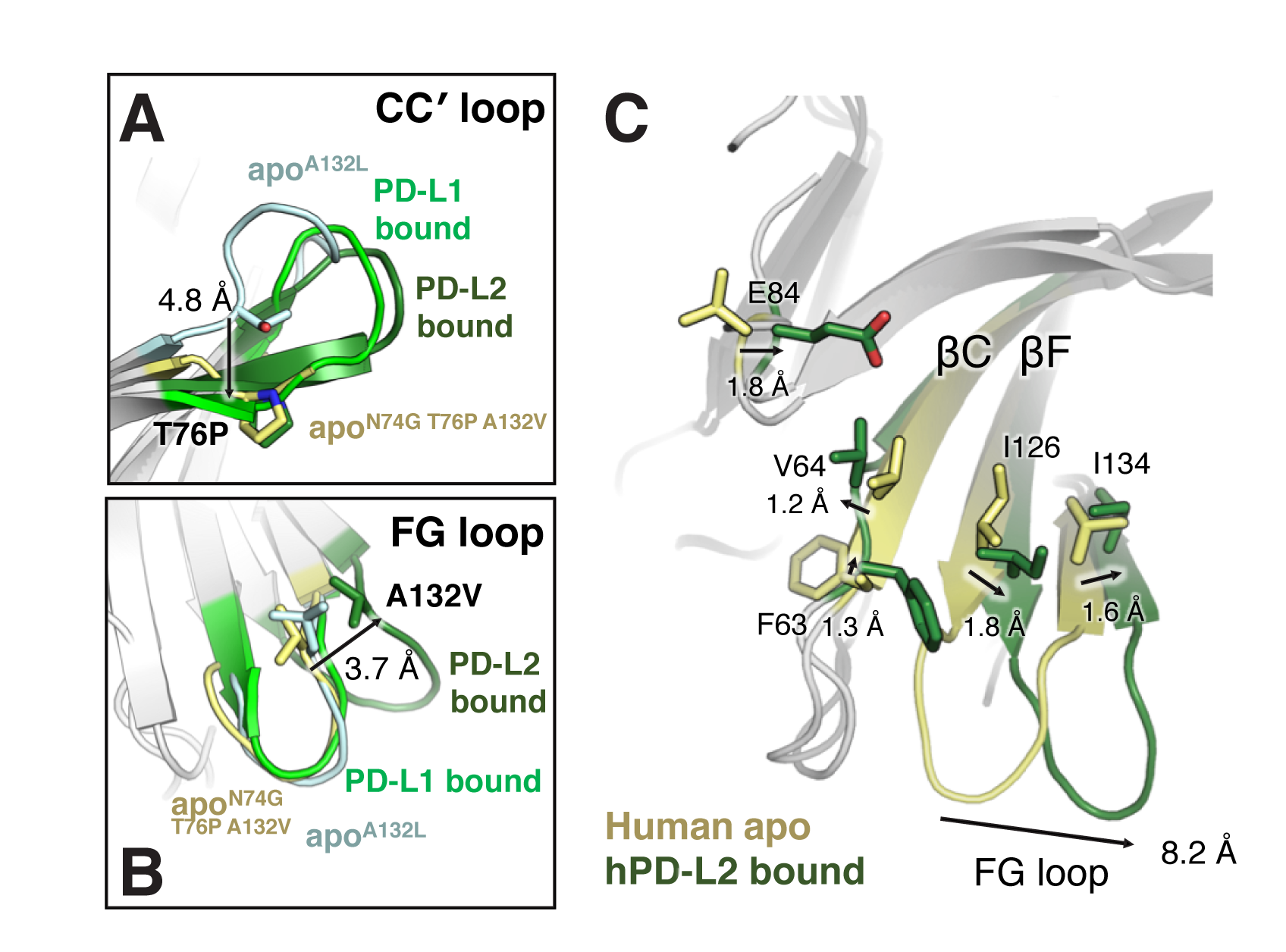
PD-L2 binding induces conformational changes in the CC’ and FG loops. (A-B) The CC and FG loops change conformations. Overlays of ribbon diagrams of (A) the CC loops and (B) the FG loops from human apo-PD-1^A132L^ (PDB: 3RRQ, cyan), apo-PD-1^N74G T76P A132V^ (pale yellow), PD-L1-bound PD-1 (PDB: 4ZQK, bright green), and PD-L2-bound PD-1^N74G T76P A132V^ (dark green). (A) T76 of apo-PD-1, as well as P76 of apo-PD-1^N74G T76P A132V^ and PD-L2- bound PD-1^N74G T76P A132V^, are indicated with sticks. The arrow highlights a 4.8 Å C shift for residue 76 (T-to-P) from apo-PD-1^A132L^ to apo-PD-1^N74G T76P A132V^. (B) L132 of apo-PD-1^A132L^, as well as V132 of apo-PD-1^N74G T76P A132V^ and PD-L2-bound PD-1^N74G T76P A132V^, are indicated with sticks. The arrow highlights a 3.7 Å C shift for V132 from apo-PD-1^N74G T76P A132V^ to PD-L2-bound PD-1^N74G T76P A132V^. (C) Pocket formation is associated with the loop change. Overlay of ribbon diagrams of human apo-PD-1^N74G T76P A132V^ (pale yellow) and PD-L2-bound PD-1^N74G T76P A132V^ (dark green). A subset of pocket residues that undergo main-chain rearrangements (arrows) are indicated with sticks. Distances of C shifts from the unbound to the PD-L2-bound states are indicated. The FG loop shift of 8.2 Å for human PD-1 was measured using the C of P130.

In contrast, the conformations of the FG loop are the same in all three apo-PD-1 structures: one with an A132L substitution in the FG loop (31) and the triple and double mutants described here (Figure S7B). Upon binding PD-L1 (19), there are no substantial conformational changes in the FG loop (Figure 6B). There is, however, a drastic shift in the FG loop conformation upon binding PD-L2 (Figures 6B and S7B).

### Structural plasticity of the human PD-1 ligand-binding interface

To further investigate how the observed loop changes are associated with pocket formation, we superimposed the apo and PD-L2-bound structures of our human triple-mutant PD-1 (Figure 6C). Upon binding PD-L2, a large conformational change occurs in the PD-1 ligand-binding interface (Figure 6C). A three-residue shortening of βC occurs (Figure S7C), and βC and βF move apart to create a deep cleft (Figure 6C). The rearrangements in the pocket propagate to the edge of the FG loop, resulting in a remarkable 8.2 Å lateral shift (Figure 6C).

We note that the overall change is less dramatic in murine PD-1 (Figure S7D). The closed architecture of the murine pocket does not require flipping of residues E84 and F63, as seen in human PD-1, and there is no secondary structure change in βC in murine PD-1 (Figure S7E). Taken together, our results provide a structural basis for systematic rearrangements at the human PD-1 ligand-binding interface that couple pocket formation and changes in the loops of PD-1 when it binds PD-L2.

## Discussion

A prominent pocket forms in human PD-1 upon binding PD-L2. This pocket has a volume of 170 Å^3^, comparable to pockets that bind small-molecule drugs (22-25). The structure of this pocket is quite distinct from the corresponding pocket in murine PD-1 bound to PD-L2 (21).

We speculate that this pocket represents an attractive drug target. How would a pocket-binding drug bind to a flat protein surface? We conceptualize an ensemble of PD-1 conformations (Figure S8) in which the predominant species of apo-PD-1 has a flat ligand-binding surface (*K*_i_ < 1). A pocket-binding drug will stabilize the PD-1 conformation containing the pocket (*K*_iii_). Drug binding via an induced-fit mechanism (*K*_iv_ > 1) can also occur.

The human PD-1/PD-L2 structure reported here will facilitate virtual drug screening to identify potential lead compounds (e.g., (30)). Specifically, we envision a small molecule binding to PD-1 contacting all or many of the residues that form the pocket, particularly F63, V64, N66, Y68, E84, L122, G124, I126, I134, and E136 in a conformation similar to that formed in the complex with PD-L2 (Figure 5B). In addition, the structures of the indole and phenyl rings and neighboring sidechains of PD-L2 when bound to the pocket (Figure 5C) are potentially useful starting points for the design of fragment-based screening scaffolds (41, 42).

Since the PD-1 pocket is not populated substantially in the absence of PD-L2, it is not straightforward to use traditional drug-screening methods to identify small molecules that bind the pocket. Nonetheless, we speculate that conformational changes in the CC′ and FG loops and formation of pockets in the ligand-binding interface of PD-1 are thermodynamically coupled (Figure 7) and that this coupling can be used to enable drug-discovery efforts. We envision that PD-1 exists in an ensemble of conformations in the absence of ligands, populating predominantly structures that contain a flat ligand-binding face (i.e., *K*_1_ < 1). PD-1 molecules with a pre-formed pocket have a higher affinity for PD-L2 (*K*_3_ > *K*_2_). Thermodynamics dictates that *K*_1_*K*_3_ = *K*_2_*K*_4_, so *K*_4_ > *K*_1_. In this model, the PD-1 loop variants studied here increase *K*_1_, and lead to a higher proportion of apo-PD-1 in the PD-L2-bound conformation.

**Figure 7.**
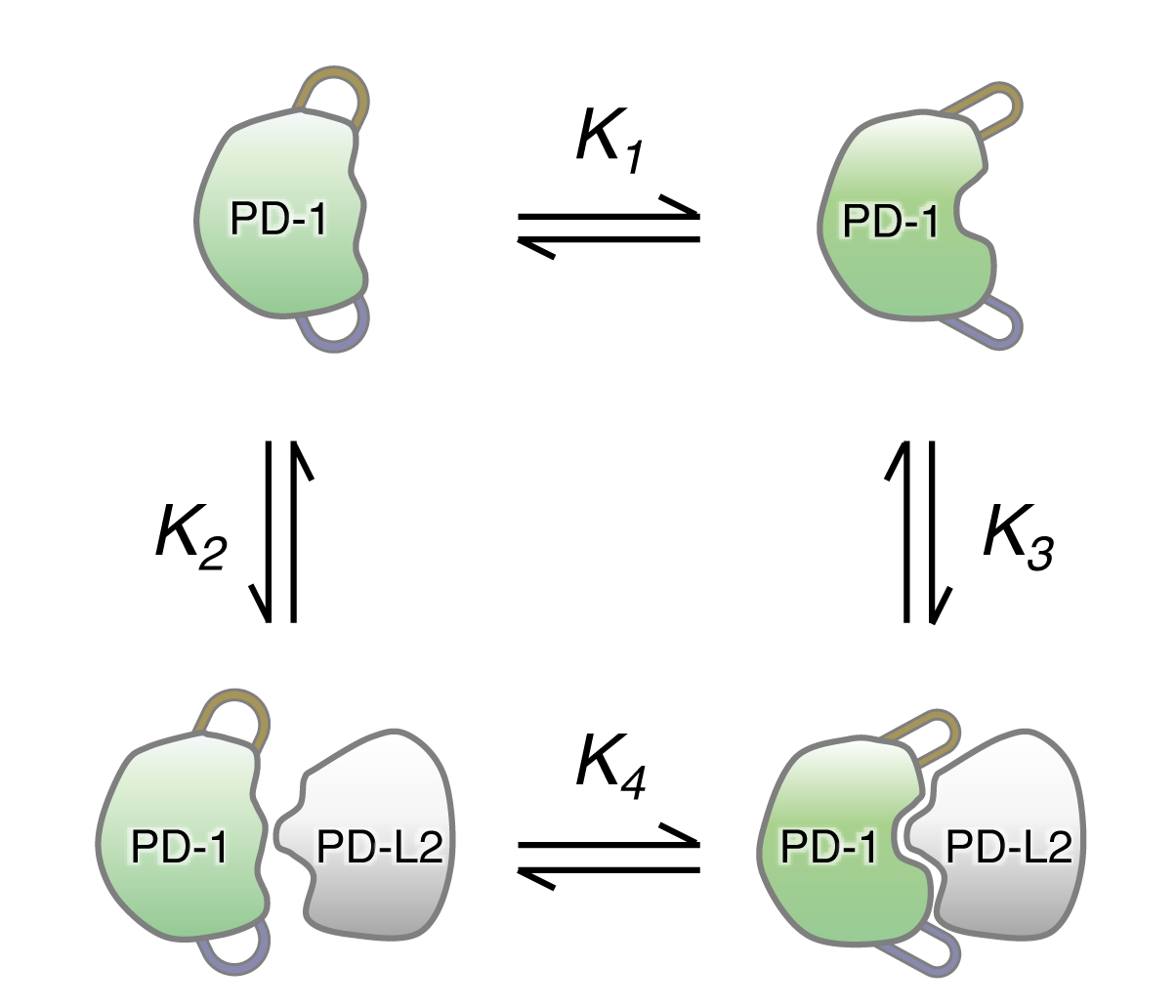
Model for coupling of pocket formation with loop movements in PD-1. A thermodynamic cycle for PD-1 binding PD-L2. For clarity, only two of the states in the conformational ensemble of apo-PD-1 (top) are depicted. In one of these states (left) the ligand-binding interface is flat. In the second state (right) a pocket is formed and the loops have moved. Mutations or external agents (e.g., antibodies) could stabilize the loops in the PD-L2 bound conformation (i.e., increase *K_1_*), thereby increasing the population of apo-PD-1 molecules in the bound conformation. Equilibrium constants for each step in the thermodynamic cycle are indicated (see Discussion).

The higher association constants (*kon*) for binding ligands by the mutant PD-1, as compared to wild-type PD-1 (Figures 3C and S2C), support this model. Such kinetic properties are consistent with an increased fraction, relative to wild-type PD-1, of unliganded mutant PD-1 molecules that are in a ligand-bound conformation ((34); see however, (35)). In addition, the CC′ loop shifts toward the PD-L2-bound conformation in the apo-PD-1 triple and double mutants (Figure S7A). (While there are only minimal changes in the pocket in both apo-PD-1 mutants (Figure 5A), the pocket residues and a neighboring FG loop have substantial crystal contacts in the lattice (Figure S5H) that likely interfere with conformational changes.)

Coupling between the pocket and the loops would stabilize the pocket in the absence of a ligand, for example if the loops were held in their PD-L2-bound conformations with antibodies or aptamers. Alternatively, or in addition, new mutations (e.g., amino acid replacements, insertions and/or deletions) could be selected for or designed to induce conformational changes in the loops. This coupling could therefore enable more traditional approaches to small-molecule drug discovery, such as high-throughput screening (43, 44) and/or the discovery of allosteric regulators of PD-1 activity. More generally, our work has implications for enhancing discovery of small-molecule inhibitors of other “undruggable” protein-protein interactions.

## Materials and Methods

Additional information is provided in the SI Appendix, SI Materials and Methods.

### Yeast surface display

Deep mutational scanning of the CC′ and FG loops of human PD-1 was performed using a previously described PCR-based method (27). PD-1 loop variant libraries were constructed using the *Saccharomyces cerevisiae* EBY100 strain (ATCC) (Table S1), by fusing PD-1 to the C-terminus of Aga2 under the *GAL1* promoter. MACS and FACS experiments were performed using recombinant human PD-L2-Fc or PD-L1-Fc proteins.

### Bio-layer interferometry

Bio-layer interferometry was performed on an Octet RED96^®^ system at 30 °C in a buffer of 150 mM NaCl, 20 mM HEPES:NaOH pH 7.4, 0.1% bovine serum albumin, and 0.05% Tween 20. Human PD-1-Fc proteins were loaded onto anti-human IgG Fc capture (AHC) biosensors, associated in defined concentrations of human PD-L2-His6 or PD-L1-His6 proteins, and then dissociated in buffer.

### Protein crystallization and X-ray crystallography

The human apo-PD-1^N74G T76P A132V^ and human apo-PD-1^T76P A132V^ proteins (Table S2) were over-expressed in and refolded from the inclusion bodies of *Escherichia coli* BL21(DE3) (Invitrogen^TM^). The human apo-PD-1^N74G T76P A132V^ protein was crystallized in 100 mM NaCl, 100 mM Tris:HCl pH 8.0, and 27% (w/v) PEG-MME 5,000. The human apo-PD-1^T76P A132V^ protein was crystallized in 100 mM NaCl, 100 mM Tris:HCl pH 8.0, and 36% (w/v) PEG 3,350. The human PD-1^N74G T76P A132V^ and human PD-L2^IgV^ protein complex (Table S2) was produced using the human Expi293F cell line (Gibco^TM^). The complex was crystallized in 200 mM magnesium acetate and 10% (w/v) PEG 8000. All X-ray diffraction data were collected at SSRL beam lines 12-2 or 14-1, and processed using HKL-3000 (45). Molecular replacement, refinement, and density modification were performed in Phenix (46) and model building in Coot (47).

### Accession number

Coordinates and structure factors for the human PD-1^N74G T76P A132V^ / PD-L2^IgV^ complex, human apo-PD-1^N74G T76P A132V^, and apo-PD-1^T76P A132V^ will be deposited in the RCSB Protein Data Bank (http://www.rcsb.org). Accession PDB IDs will be available after deposition. Structures are available immediately at https://peterkimlab.stanford.edu.

## Acknowledgments

We thank Drs. J.S. Fraser and J.S. Weissman for helpful comments on an earlier version of this manuscript. We also thank members of the Kim laboratory, especially B.N. Bell, T.U.J. Bruun, M.V.F. Interrante, P.A. Weidenbacher, Drs. L.N. Deis, Y. Hwang Fu, L.W.H. Lee and A.E. Powell for discussion and helpful comments on the manuscript, Drs. J.S. Fraser, J.D. Bloom and L. Zhang for insightful discussion and technical expertise. We thank Dr. D. Fernandez of Stanford ChEM-H Macromolecular Structure Knowledge Center (MSKC), and staff scientists of the Stanford Synchrotron Radiation Lightsource (SSRL) beam lines 12-2 and 14-1 for X-ray crystallographic data collection. Use of the SSRL, SLAC National Accelerator Laboratory, is supported by the U.S. Department of Energy, Office of Science, Office of Basic Energy Sciences under Contract No. DE-AC02-76SF00515. The SSRL Structural Molecular Biology Program is supported by the DOE Office of Biological and Environmental Research, and by the NIH NIGMS P41GM103393. This work was supported by the Emerson Collective Cancer Research Fund, NIH DP1 DA043893, the Virginia and D. K. Ludwig Fund for Cancer Research, and the Chan Zuckerberg Biohub. S.T. is a Merck Fellow of the Damon Runyon Cancer Research Foundation, DRG-2301-17.

## Supplementary Information for

A high-affinity human PD-1/PD-L2 complex informs avenues for small-molecule immune checkpoint drug discovery

Shaogeng Tang and Peter S. Kim

## SI Materials and Methods

### Deep mutational scanning

Deep mutational scanning of the CC′ and FG loops was performed using a previously described PCR-based method (1). For the amplicon, we used the cDNA encoding human PD-121-150 with 120 base-pairs of upstream and 227 base-pairs of downstream DNA sequences in the yeast-display plasmid of pST892 - pRS414 PGAL1- AGA2-PD121-150-Myc. The upstream sequence is from the Factor Xa site to PD-1P20, and the downstream is from PD-1E150 to the end of the ADH1 terminator. This amplicon was produced by PCR and diluted to 3 ng/μL. We combined forward oligonucleotides that contained a randomized NNN trinucleotide in codons encoding amino acids of the CC′ (70MSPSNQTDK78) and/or FG (127SLAPKAQ133) loops of human PD-1 in equimolar quantities to create three forward-mutagenesis primer pools: one for the CC′ loop, one for the FG loop, and one combining both loops. We also combined the reverse complement of each of these oligonucleotides in equimolar quantities to create three reverse-mutagenesis primer pools. The Tm of all primers were designed to be ∼59-62 oC.

Two fragment PCR reactions were set up for each template. The forward-fragment reactions contained 15 μL of CloneAmp HiFi PCR premix (Takara Bio USA, Inc), 2 μL of the forward mutagenesis primer pool at a total oligonucleotide concentration of 4.5 μM, 2 μL of 4.5 μM OST939 ADH1 terminator reverse primer and 3 ng/μL of the aforementioned PD-1 amplicon as a template, and 7 μL of ddH2O. The reverse-fragment reactions were identical, except that the reverse mutagenesis pool was substituted for the forward mutagenesis pool, and OST939 was substituted for OST938 Factor Xa forward primer. PCRs were performed using the following program: (1) 98 oC, 2 min, (2) 98 oC, 10 sec, (3) 72 oC, 1 sec, (4) 65 oC cooling to 50 oC at 0.5 oC/sec, (5) go to step (2), 6 times, (6) 4 oC, hold.

The PCR products were diluted 1:4 in ddH2O and used for stitching PCR reactions: 15 μL CloneAmpTM HiFi PCR premix, 4 μL of the 1:4 dilution of the forward-fragment reaction, 4 μL of the 1:4 dilution of the reverse-fragment reaction, 2 μL of 4.5 μM OST938, 2 μL of 4.5 μM OST939 and 3 μL of ddH2O. The stitching PCRs were performed using the same PCR program except with 20 thermal cycles. The PCR products were purified over 1% agarose-TAE gels. Purified products were subsequently used as templates for a second round of fragment reactions followed by another stitching PCR. The second-generation PD-1 loop variant libraries were generated using the same method, except using either the cDNA of PD-1T76P or PD-1A132V as an amplicon, and re-designed oligonucleotide pools containing either T76P or A132V substitution for fragment PCRs.

The vector backbone was generated by PCR to linearized pST892 using an OST941 PD1P21 reverse primer and an OST880 PD-1E150 forward primer. 16 μg of purified inserts were combined with 4 μg of purified vector backbone. The DNA mixtures were precipitated at −80 oC for 2 hours after mixing with 1:10 volume of 3.0 M NaOAc and three-time volume of 100% EtOH. The DNA pellet was collected by centrifugation at 21,300 rcf for 30 min at 4 oC, and then air-dried.

#### Assembly of the PD-1 loop variant libraries

The PD-1 variant libraries of the CC′ loop, the FG loop, and the CC′ and FG loops were generated separately. Saccharomyces cerevisiae EBY100 strain (ATCC) (Table S1) was inoculated in 5 mL of YPD media overnight. The stationary-phase culture was diluted to OD600 of 0.1 in 50 mL YPD. After reaching mid-log phase, 500 μL of 1 M Tris:HCl pH 8.0 and 2.5 M dithiothreitol was added to the culture for 15 min. Yeast cells were then collected and washed twice with a buffer of 10 mM Tris:HCl pH 7.5, 270 mM sucrose, 1 mM MgCl2, and resuspended in a final volume of 300 μL. The air-dried insert-and-vector DNA mixture pellets were mixed with the cells and aliquoted into 50-100 μL 0.2 cm gap Gene Pulser®/MicroPulser™ electroporation cuvettes (BioRad). Yeast transformations were performed by electroporation at 0.54 kV and 25 μF. Yeast cells were recovered in 10 mL YPD media at 30 oC for 1 hour, and then resuspended in 50 mL SD-CAA media supplemented with 10% penicillin-streptomycin and incubated on a shaking platform at 30 oC. At day 2, ∼50 OD600V of yeast cells were passaged into 100 mL SD-CAA media and incubated at 30 oC for one day.

#### Selection of yeast libraries by MACS and FACS

The PD-1 variant libraries of the CC′ loop, the FG loop, and the CC′ and FG loops were induced in 200 mL of SG-CAA media with 10% penicillin-streptomycin at 20 oC for 48 hours. For selection round 1 using MACS, cell pellets of ∼109 yeast of each library were combined, washed with 25 mL, and resuspended in 5 mL of a selection buffer of 150 mM NaCl, 20 mM HEPES:NaOH pH 7.4, 0.1% Bovine Serum Albumin (Sigma-Aldrich) and 2.5 mM EDTA:HCl pH 8.0. For negative selection, yeast were incubated with 1 μM recombinantly purified human Fc pre-mixed with 100 μL of anti-human IgG microbeads (Miltenyi Biotec) at 4 oC for 1 hour, and then loaded onto a LD column (Miltenyi Biotec) attached to a MACS magnetic MultiStand (Miltenyi Biotec). The LD column was washed with 10mL of selection buffer. Flow-through was collected and resuspended in 5 mL. For positive selection, these cells were then incubated with 500 nM PD-L2-Fc pre-mixed with 100 μL of anti-human IgG microbeads at 4 oC for 3 hours, and then loaded onto a new LD column. The column was washed with 20 mL of selection buffer and then removed from the magnetic MultiStand. Cells were eluted by gently plunging the LD column with 10 mL of selection buffer and then resuspended in 10 mL SD-CAA with 10% penicillin-streptomycin. After growing at 30 oC for 24 hours, the cells were induced in SG-CAA media at 20 oC for 48 hours.

For selection round 2 using FACS, 1 mL of induced cells from round 1 at stationary-phase were incubated with 1 nM PD-L2-Fc at 4 oC for 2 hours and then washed and resuspended in 1 mL of the selection buffer. The cells were incubated with 1:250 dilution of the anti-human Fc-Alexa Fluor 647 antibody (BioLegend) and 1:250 dilution of the anti-cMyc-Alexa Fluor 488 antibody (Novus Biologicals) at 4 oC for 30 min, washed twice and resuspended in 1 mL of selection buffer. FACS was performed with a SH800S cell sorter (SONY). Using unlabeled yeast as a reference, PD-L2-Fc (Alexa Fluor 647 labeled) positive yeast cells were gated and sorted in 3 mL SD-CAA media. Sorted cells were grown at 30 oC for 48 hours, and then induced in SG-CAA media at 20 oC for 24 hours.

Selection for enhanced PD-L2 binding using a second-generation library of PD- 1T76P loop variants was performed with the same method, except for labelling with 50 nM PD-L2-Fc in round 1 of MACS, and 100 pM PD-L2-Fc in round 2 of FACS.

Selection for enhanced PD-L1 binding using the first-generation PD-1 loop variant library was performed with the same method. Cells were labeled with 1000 nM PDL1-Fc for round 1 of MACS and 500 pM PD-L1-Fc for round 2 of FACS. The second-generation PD-1A132V loop variant library was selected with 100 nM PD-L1-Fc for round 1 of MACS and 100 pM PD-L1-Fc for round 2 of FACS.

#### Flow Cytometry

For yeast staining, polyclonal (a library) or monoclonal (a single-plasmid transformation) yeast were inoculated in SD-CAA at 30 oC to reach stationary phase, and induced in SG-CAA media at 20 oC for 24-48 hours. ∼1 OD600V cells were pelleted, washed and resuspended in ∼50 μL. 1 μL of resuspended yeast were incubated for 45 min with 100 μL recombinant bait proteins (e.g.: PD-L2-Fc or PD-L1-Fc) that were diluted with the selection buffer of 150 mM NaCl, 20 mM HEPES:NaOH pH 7.4, 0.1% BSA and 2.5 mM EDTA:HCl pH 8.0 into desired concentrations. Yeast cells were then washed with 1 mL of selection buffer and resuspended in 100 μL of selection buffer with the addition of 0.5 μL of anti-human Fc-Alexa Fluor 647 antibody, and 0.5 μL of dilution of the anti-cMyc-Alexa Fluor 488 or anti-cMyc-FITC antibody (Miltenyi Biotec) for 15 min. Yeast cells were then washed with 1 mL and resuspended in 100 μL of the selection buffer. Flow cytometry was performed using a BD Accuri™ C6 Flow cytometer. 10,000-100,000 events were recorded and analyzed for clonal expression (the FL1 channel) and bait binding (the FL4 channel). Graphs of flow cytometry were generated using FlowJo.

#### Protein expression and purification

Recombinant PD-1, PD-L1, and PD-L2 proteins used for bio-layer interferometry were produced in Expi293F cells. Supernatants of the cell culture were collected for protein purification. Fc-fusion proteins (PD-1-Fc, PD-L1-Fc and PD-L2-Fc) were purified by PierceTM protein A agarose (Thermo Fisher Scientific). The resins were batch bound, washed with 150 mM NaCl, 20 mM HEPES:NaOH pH 7.4 and eluted with 100 mM glycine:HCl pH 2.8 directly into 1:10 volume of 1.0 M HEPES:NaOH pH 7.4. Supernatants of cell culture containing His6-tagged proteins (PD-L1-His6 and PD-L2- His6) were dialyzed in 150 mM NaCl, 20 mM HEPES:NaOH pH 7.4 and purified by HisPurTM Ni-NTA resins (Thermo Fisher Scientific). The resins were batch bound, washed with 150 mM NaCl, 20 mM HEPES:NaOH pH 7.4, 20 mM imidazole, and eluted with 150 mM NaCl, 20 mM HEPES:NaOH pH 7.4, 400 mM imidazole. After affinity purification, all proteins were subsequently purified by gel filtration using an ÄKTATM pure chromatography system with a Superdex 200 Increase 10/300 GL column (GE Healthcare) in 150 mM NaCl, 20 mM HEPES:NaOH pH 7.4.

#### Bio-layer interferometry

Bio-layer interferometry was performed on an Octet RED96® system (Pall ForteBio) in a buffer of 150 mM NaCl, 20 mM HEPES:NaOH pH 7.4, 0.1% BSA and 0.05% Tween 20 at 30 oC at a shaking speed of 1,000. The human PD-1-Fc proteins were loaded onto anti-human IgG Fc capture (AHC) biosensors with a load threshold of 0.8 nanometer. After loading, the biosensors were baselined, and associated in defined concentrations of human PD-L1-His6 or PD-L2-His6 proteins for 120 seconds, and then dissociated in buffer for 120 seconds. The baseline-corrected binding curves were analyzed with GraphPad Prism 8.

#### Protein purification for crystallization

For the human apo-PD-1N74G T76P A132V and human apo-PD-1T76P A132V proteins, cDNA encoding PD-1 residues 33-150 with substitutions of C93S N74G T76P A132V or C93S T76P A132V were subcloned into a pET23d vector. The C-terminus of PD-1 is fused with a GS linker and followed by a Strep-tag®II (WSHPQFEK) for affinity purification (Table S2). The recombinant proteins were overexpressed in inclusion bodies of Escherichia coli BL21(DE3) cells, with 1 mM IPTG at 18 oC overnight. Cell pellets were collected and sonicated in 500 mM NaCl, 20 mM HEPES:NaOH pH 7.4. The inclusion bodies were obtained by centrifugation at 15,000 rcf for 45 min at 4 oC, and solubilized in 500 mM NaCl, 20 mM HEPES:NaOH pH 7.4, 8 M urea. After another round of centrifugation at 15,000 rcf for 45 min at 4 oC, the solubilized inclusion bodies were drop-by-drop mixed with 5- to 10-fold volume of a refolding buffer of 24 mM NaCl, 1 mM KCl, 50 mM HEPES:NaOH pH 7.4, 500 mM L-arginine, 9 mM glutathione and 1 mM glutathione disulfide. The refolded mixture was cleared by centrifugation at 15,000 rcf for 45 min at 4 oC and passed through a 0.22 μm filter, and then batch bound with Strep-Tactin®XT Superflow® high capacity resins (IBA Lifesciences). The resins were washed with 150 mM NaCl, 100 mM Tris:HCl pH 8.0, 1 mM EDTA, and eluted with 150 mM NaCl, 100 mM Tris:HCl pH 8.0, 1 mM EDTA, 50 mM biotin. The PD-1 proteins were further purified by gel filtration using an ÄKTATM pure chromatography system with a Superdex 200 Increase 10/300 GL column in 150 mM NaCl, 20 mM HEPES:NaOH pH 7.4.

For the human PD-1N74G T76P A132V and human PD-L2IgV protein complex, human cDNA encoding PD-1 residues 33-150 with substitutions of C93S N74G T76P A132V and N-linked glycosylation mutations of N49S N58S N74D N116D, and human cDNA encoding PD-L2 residues 1-123 with N-linked glycosylation mutations of N37D N64D were separately subcloned into pADD2 vectors. The N-terminus of the PD-1 protein was fused to a signal sequence of MGWSCIILFLVATATGVHS before residue 33. The C-terminus was fused to a PEA tag after amino acid E150, which together consisted of a C-tag for affinity purification (Table S2). Both plasmids were combined at a 1:1 ratio and used for co-transfecting Expi293F cells. Supernatants of the cell culture were collected and incubated with CaptureSelect C-tagXL affinity matrix (Thermo Fisher Scientific). The resins were washed with 150 mM NaCl, 20 mM HEPES:NaOH pH 7.4, and eluted with 2 M MgCl2, 20 mM Tris:HCl pH 8.0. The complex was further purified by gel filtration using an ÄKTATM pure chromatography system with a Superdex 200 Increase 10/300 GL column in 150 mM NaCl, 20 mM HEPES:NaOH pH 7.4. Fractions corresponding to the PD-1/PD-L2 complex were analyzed by SDS-PAGE and used for protein crystallization.

#### Protein crystallization

The human apo-PD-1^N74G T76P A132V^ protein was crystallized at room temperature in a hanging-drop vapor diffusion system by mixing 1 μL of 9.8 mg/mL of protein with 1 μL of 1 mL reservoir solution containing 100 mM NaCl, 100 mM Tris:HCl pH 8.0, 27 % (w/v) PEG-MME 5,000. Crystals were transferred into the same solution supplemented with 20.5% (w/v) PEG 200 before cooling to liquid nitrogen temperature.

The human apo-PD-1^T76P A132V^ protein was crystallized at room temperature in a hanging-drop vapor diffusion system by mixing 1 μL of 8.4 mg/mL of protein with 1 μL of 1 mL reservoir solution containing 100 mM NaCl, 100 mM Tris:HCl pH 8.0, 36% (w/v) PEG 3,350. Crystals were transferred into the same solution supplemented with 12% (w/v) PEG 200 before cooling to liquid nitrogen temperature.

The human PD-1^N74G T76P A132V^ and human PD-L2^IgV^ protein complex was crystallized at room temperature in a sitting-drop vapor diffusion system by mixing 200 nL of 12.6 mg/mL of proteins with 200 nL of 80 μL reservoir solution containing 200 mM magnesium acetate, 10% (w/v) PEG 8000 (Anatrace MCSG4 H12). Crystals were transferred into the same solution supplemented with 28% (w/v) PEG 8,000 before cooling to liquid nitrogen temperature.

#### X-ray crystallography

For the human apo-PD-1^N74G T76P A132V^ crystal, X-ray diffraction data was collected to 1.18 Å at the Stanford Synchrotron Radiation Lightsource (SSRL) beam line 12-2 of SLAC National Accelerator Laboratory. The crystal belonged to the space group P 32 2 1 with unit cell dimensions a=46.2 Å b=46.2 Å c=89.3 Å α=90° β=90° γ=120° (Table 1).

For the human apo-PD-1T76P A132V crystal, X-ray diffraction data was collected to 1.42 Å at the SSRL beam line 14-1. The crystal also belonged to space group P 32 2 1 with unit cell dimensions a=46.2 Å b=46.2 Å c=89.4 Å α=90° β=90° γ=120° (Table 1).

For the human PD-1^N74G T76P A132V^/PD-L2^IgV^ cocrystal, X-ray diffraction data was collected to 1.99 Å at the SSRL beam line 12-2. The crystal belonged to space group P 2_1_ 2_1_ 2_1_ with unit cell dimensions a=41.3 Å b=67.8 Å c=89.7 Å α=90° β=90° γ=90° (Table 1).

All diffraction data were processed using HKL-3000 (2). The structure of human apo-PD-1^N74G T76P A132V^ was solved using Phaser in Phenix (3) by molecular replacement with the human apo-PD-1^A132L^ structure (PDB: 3RRQ) as a search model. The structure of human apo-PD-1^T76P A132V^ was solved by molecular replacement with the human apo-PD-1N74G T76P A132V structure. The structure of the human PD-1^N74G T76P A132V^/PD-L2^IgV^ complex was solved by molecular replacement with the human apo-PD-1N74G T76P A132V structure, and the IgV domain of murine PD-L2 from the murine PD-1/PD-L2 structure (PDB: 3BP5). Model refinement and density modification were performed in Phenix (3). Model building was performed using Coot (4). Structural images were generated with PyMOL (Schrödinger). Pocket volumes and volume-depths were measured using POCASA 1.1 (5), with parameter settings of 6 Å for probe radius, 16 for single point flag, 18 for protein-depth flag and 1.0 Å for grid size.

**Figure S1.**
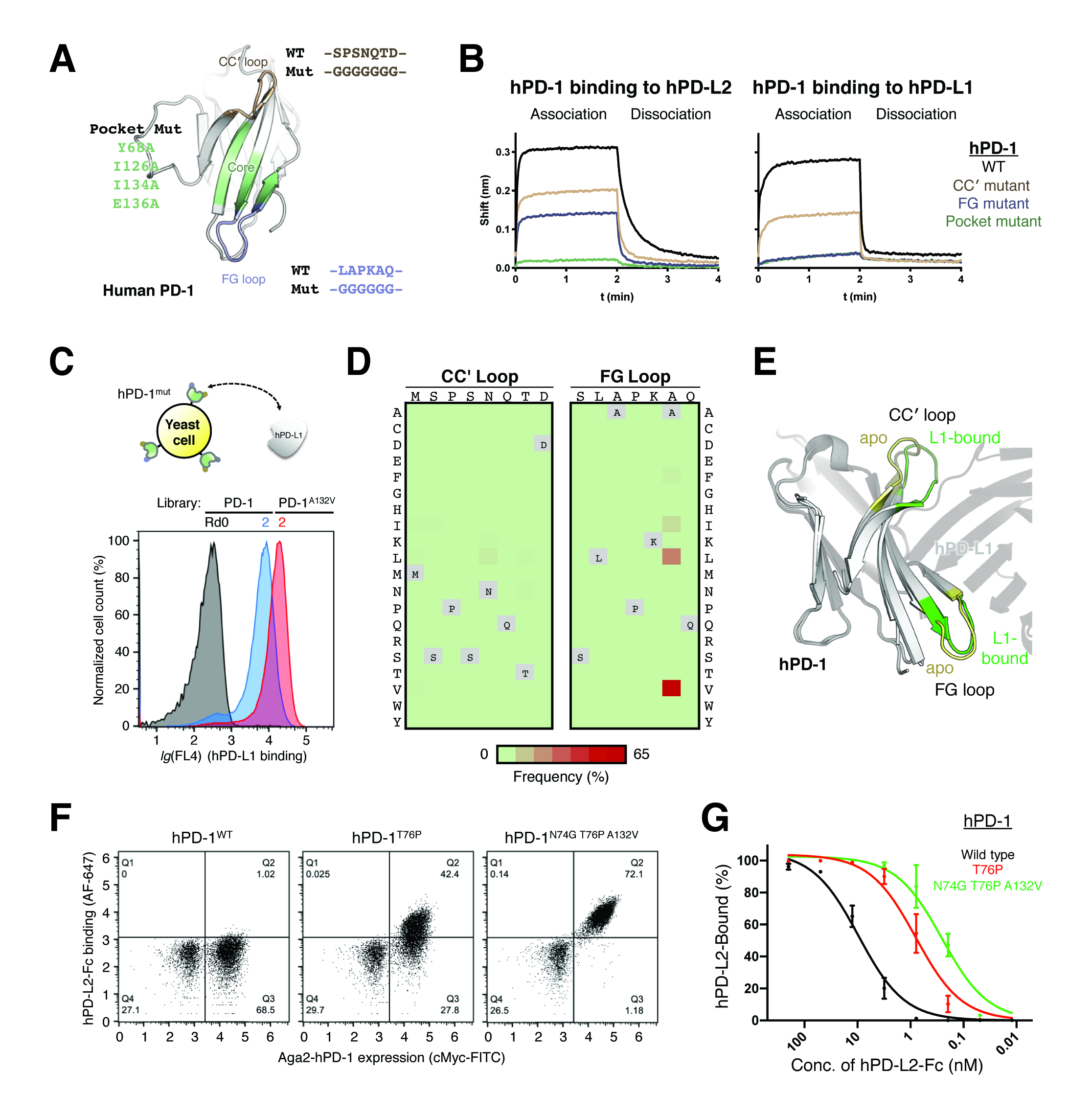
Engineering PD-1 loop variants with enhanced PD-L2 affinity, related to Figure 2. (A) Ribbon diagram of the human PD-1 ectodomain, with the CC loop colored in wheat, the FG loop in light blue, and the hydrophobic core/pocket in pale green. Glycine mutants were designed for the CC loop (residues 71-77) and the FG loop (residues 128-133), and an alanine mutant for the pocket (Y68A I126A I134A E136A). WT, wildtype; Mut, mutant. (B) Bio-layer interferometry of the binding of sensor-loaded PD-1, the glycine-loop mutants, and the pocket mutant to 1.9 M PD-L2 (left) and 17 M PD-L1 (right). Corresponding PD-1-Fc proteins were loaded onto anti-human IgG Fc capture (AHC) biosensors. Association and dissociation were each monitored for 2 min. (C) Schematic of yeast surface display of a human PD-1 loop variant library and selection with a recombinant human PD-L1 ectodomain. Overlay of flow-cytometric histograms of the PD-1 loop variant yeast library at selection rounds 0 (black), 2 (blue), and the PD-1^A132V^ loop variant yeast library at selection round 2 (red). Yeast cells were stained with 100 nM PD-L1-Fc, followed by Alexa Fluor 647-labeled secondary antibody against human Fc. (D) Frequency heatmaps of the amino acid substitutions of PD-1 in the CC loop (left) and the FG loop (right) after selection round 2 of the PD-1 loop variant yeast library using PD-L1-Fc. (E) Overlay of ribbon diagrams of human apo-PD-1^A132L^ (PDB: 3RRQ) and human PD-L1-bound human PD-1 (PDB: 4ZQK). The CC loops and the FG loops are highlighted for the apo structure (pale yellow) and the PD-L1-bound structure (bright green). (F) Yeast-displayed PD-1^WT^ (left), PD-1^T76P^ (middle), and PD-1^N74G T76P A132V^ (right) stained with 780 nM PD-L2-Fc, followed by Alexa Fluor 647-labeled secondary antibody against human Fc. Expression of PD-1-cMyc on the yeast surface was monitored with a fluorescein isothiocyanate (FITC)-labeled cMyc antibody. (G) Titration curves of yeast-displayed PD-1^WT^ (black), PD-1^T76P^ (red), and PD-1^N74G T76P A132V^ (green) stained with PD-L2-Fc, followed by Alexa Fluor 647-labeled secondary antibody against human Fc. Fitting was performed in Graphpad Prism 8 using built-in equations of “One site - Specific binding”. Means and standard deviations were calculated from three independent experiments.

**Figure S2.**
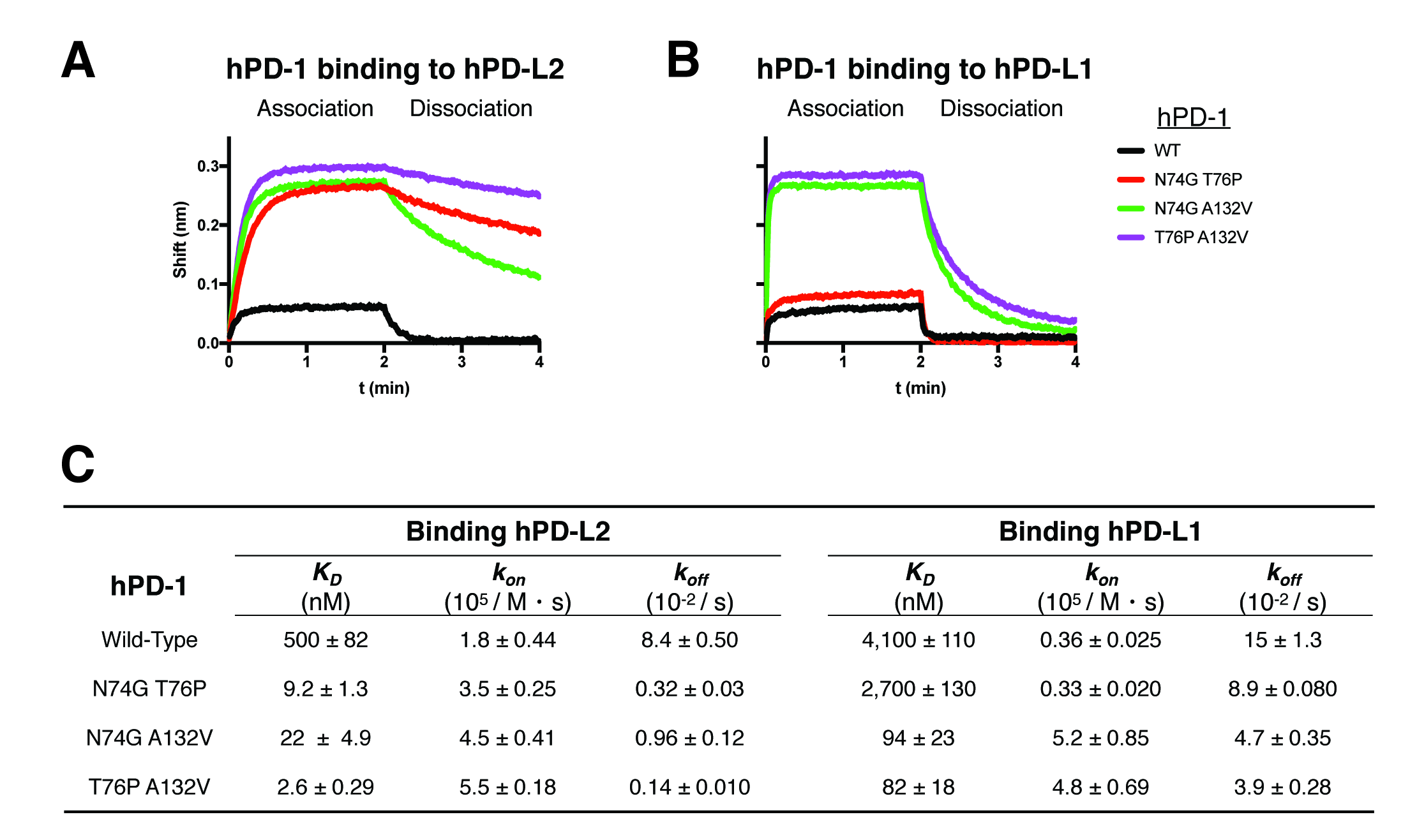
PD-1 loop variants with increased affinity and association kinetics to PD-L2 and PD-L1, related to Figure 3. (A-B) Bio-layer interferometry of the binding of sensor-loaded PD-1 and the loop variants to (A) 190 nM PD-L2 and (B) 1.1 M PD-L1. Corresponding PD-1-Fc proteins were loaded onto anti-human IgG Fc capture (AHC) biosensors. Association and dissociation were each monitored for 2 min. (C) Summary of binding affinity (KD) and kinetic parameters (association constant, kon; dissociation constant, koff) for the PD-1 loop variants binding to PD-L2 or PD-L1. Fitting of binding curves was performed in Graphpad Prism 8 using built-in equations of “Receptor binding - kinetics” models. Means and standard deviations were calculated from 3-4 independent experiments. The wild-type data in Figure 3 are presented here for comparison.

**Figure S3.**
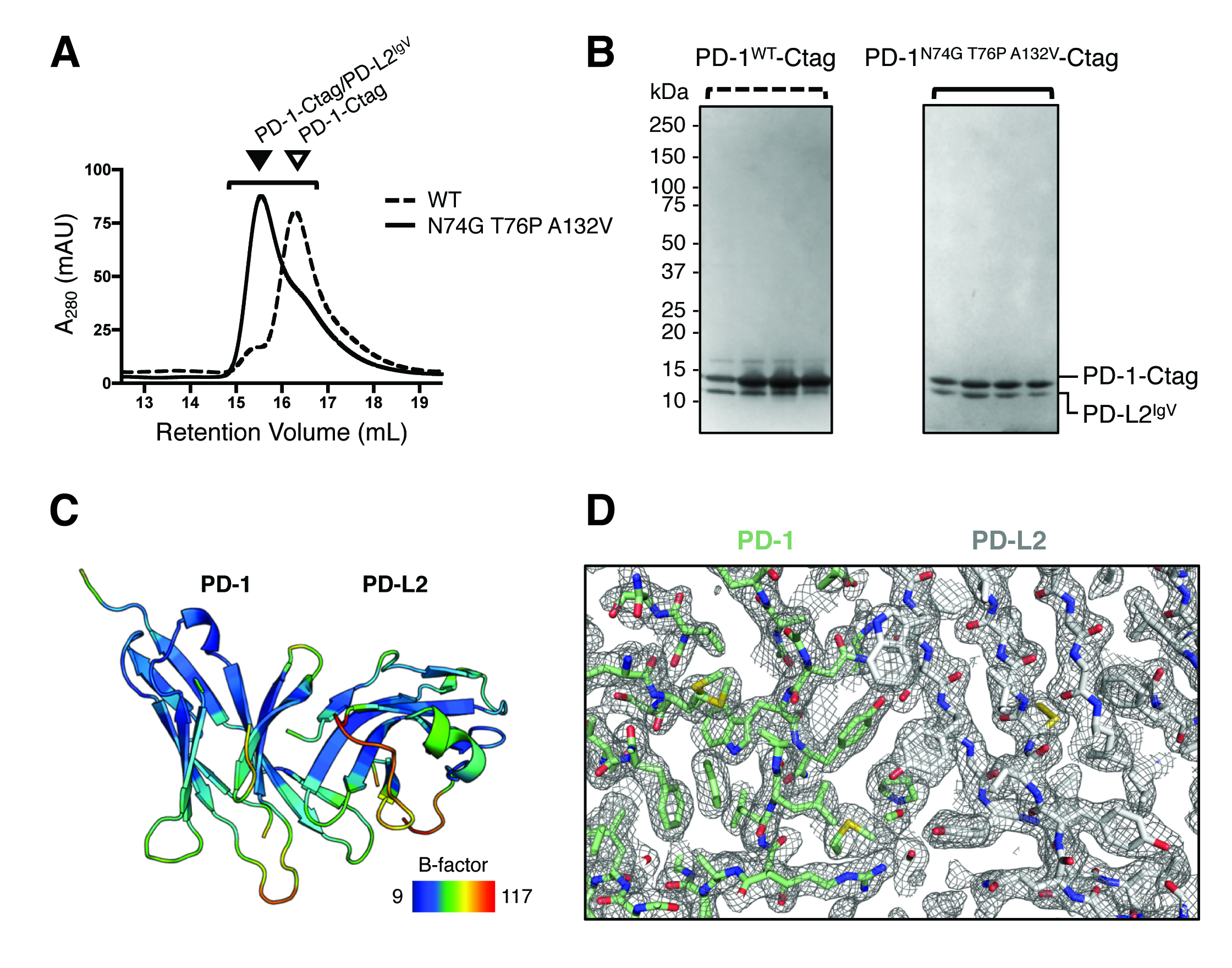
Purification and X-ray crystallography of the human PD-1/PD-L2 complex, related to Figure 4. (A) Chromatograms of size exclusion of the PD-1^WT^-Ctag/PD-L2^IgV^ complex (dashed line) and the PD-1^N74G T76P A132V^-Ctag/PD-L2^IgV^ complex (solid line). Triangles indicate the PD-1/PD-L2^IgV^ complex (left) or free PD-1 (right) proteins. The bracket indicates the fractions of both preps used for SDS-PAGE shown in (B). WT, wildtype; A_280_, absorbance at 280 nm; mAU, milli absorbance unit. (B) SDS-PAGE of fractions from size-exclusion chromatography of PD-1^WT^-Ctag/PD-L2^IgV^ (left) and PD-1^N74G T76P A132V^-Ctag/PD-L2^IgV^ (right) stained with GelCode blue safe^®^ dye. (C) A ribbon diagram displaying B-factors of the structure of the human PD-1^N74G T76P A132V^/PD-L2^IgV^ complex. (D) *2F_obs_-F_calc_*simulated-annealing composite-omit (5% atoms) electron density map contoured at 1.0 of the human PD-1^N74G T76P A132V^/PD-L2^IgV^ complex.

**Figure S4.**
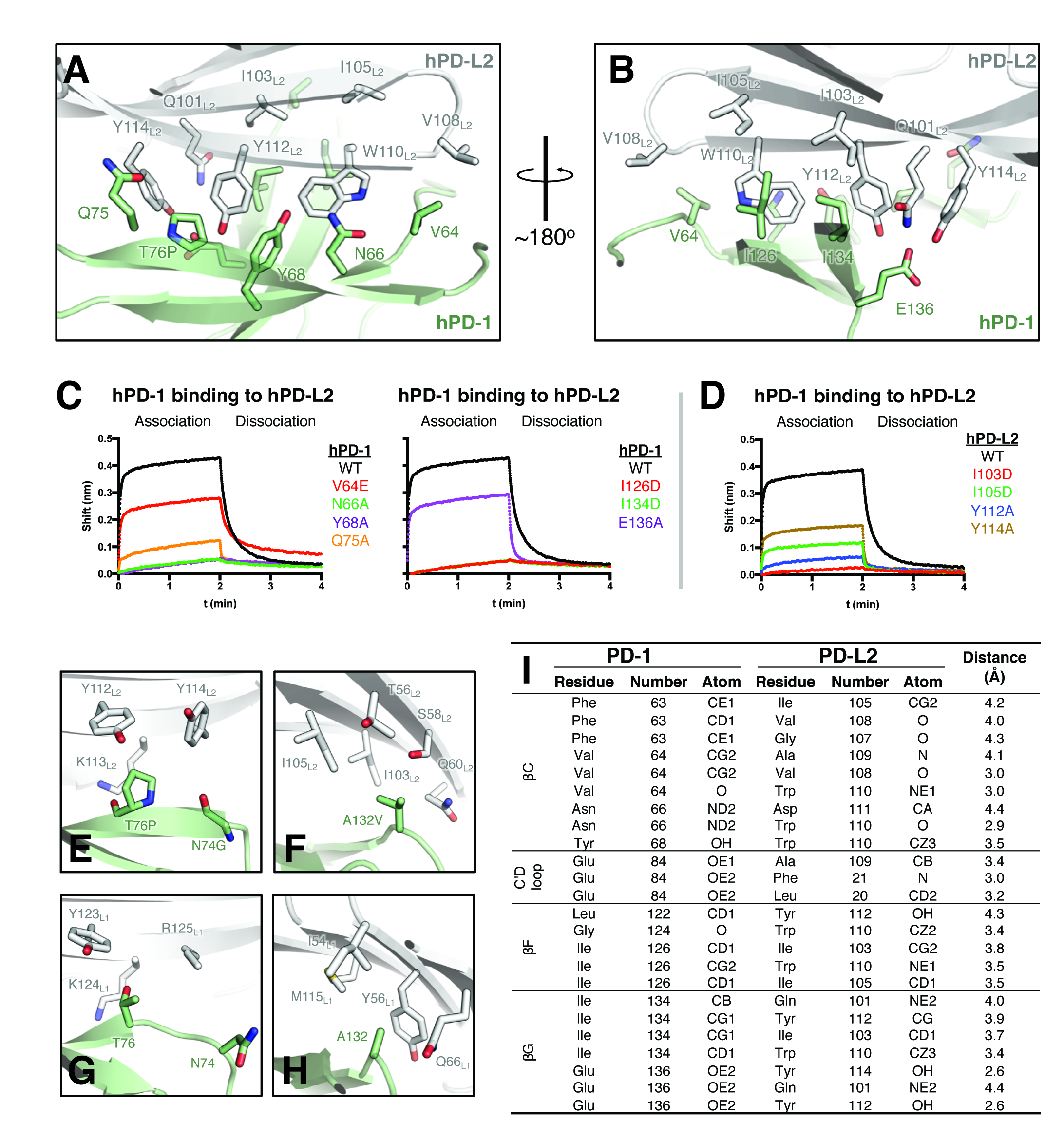
The human PD-1/PD-L2 binding interface, related to Figures 4 and 5. (A-B) Ribbon diagrams of the interface between human PD-1 (pale green) and PD-L2 (grey), overlaid with interacting residues shown as sticks. There is a ∼180o rotation between the views in (A) and (B). (C) Bio-layer interferometry of the binding of sensor-loaded PD-1 and PD-1 interface mutants to 4 M PD-L2. Corresponding PD-1-Fc proteins were loaded onto anti-human IgG Fc capture (AHC) biosensors. Association and dissociation were each monitored for 2 min. The same wild-type (WT) data are plotted for comparison. (D) Bio-layer interferometry of the binding of sensor-loaded PD-1 to 4 M PD-L2 or the PD-L2 interface mutants. WT PD-1-Fc proteins were loaded onto anti-human IgG Fc capture (AHC) biosensors. Association and dissociation were each monitored for 2 min. (E-F) Close-up ribbon diagrams of the localizations of the loop substitutions overlaid as sticks of (E) the mutated G74 and P76, and (F) V132 residues in the structure of human PD-1 (pale green) and PD-L2 (grey). PD-L2 residues are overlaid as sticks and labeled with an L2 subscript. P76 of the CC loop of PD-1 localizes between sidechains of Y112L2 and Y114L2. V132 of the FG loop localizes to a groove of T56L2, S58L2, I103L2, and I105L2. (G-H) Close-up ribbon diagrams of the localizations of the loop substitutions (overlaid as sticks) of (G) N74 and T76, and (H) A132 in the structure of human PD-1 (pale green) and PD-L1 (grey) (PDB: 4ZQK). PD-L1 residues are overlaid as sticks and labeled with an L1 subscript. Compared to PD- L2, the corresponding Y114L2 is substituted by R125L1. A132 of the FG loop localizes to a groove of I54L1, Y56L1, Q66L1, and M115L1 in PD-L1. (I) List of atoms from PD-L2 residues within 4.5 Å distance of the PD-1 pocket residues in Figure 5B. Distance measurements were generated by COCOMAPS.

**Figure S5.**
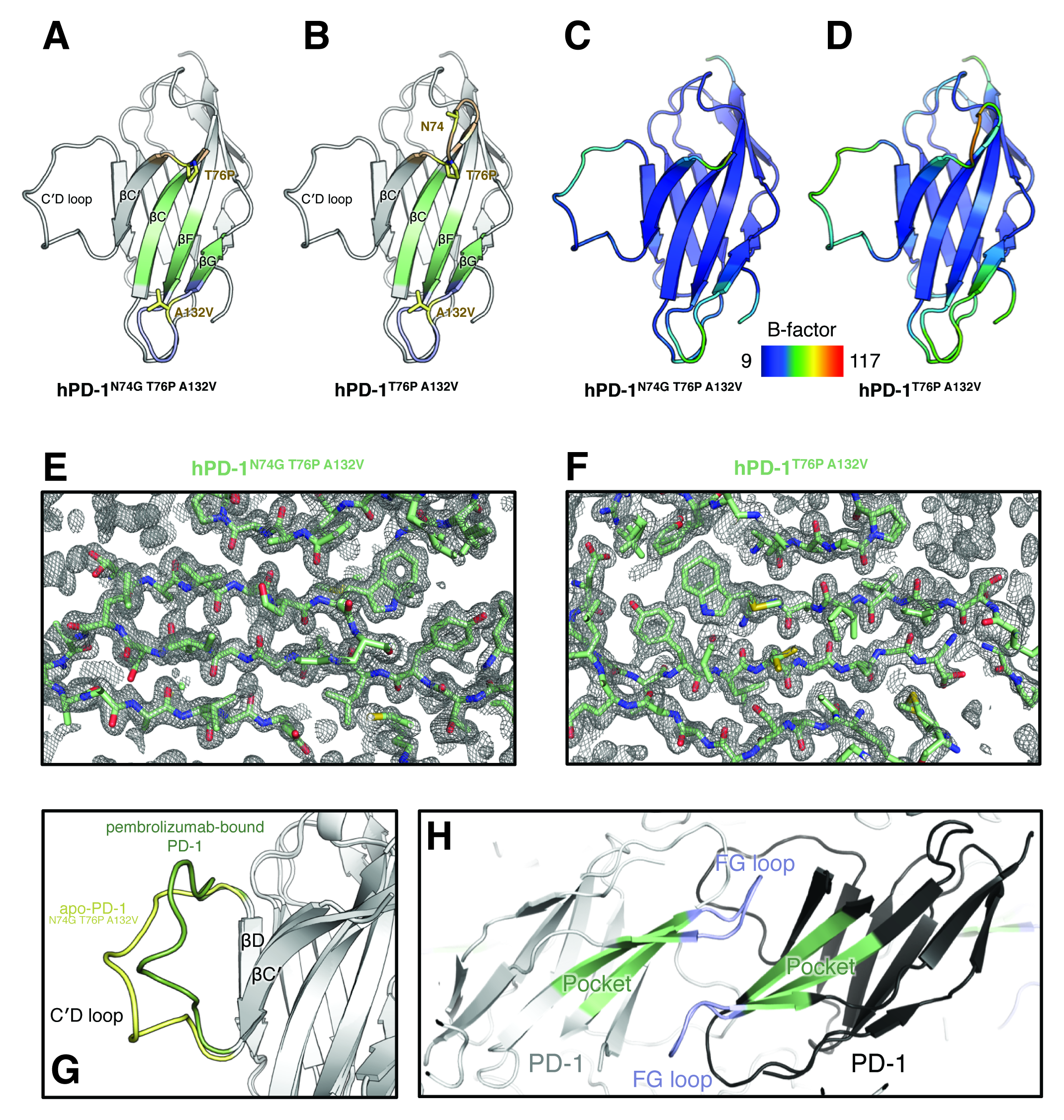
X-ray crystal structures of human apo-PD-1 loop variants, related to Figure 5. (A-B) Ribbon diagrams of the front face of (A) a 1.2 Å resolution structure of human apo-PD-1N74G T76P A132V and (B) a 1.4 Å resolution structure of human apo-PD-1T76P A132V. The CC loop (wheat), the FG loop (light blue), and the hydrophobic core/pocket (pale green) are shown. Sidechains of the loop substitutions appear as sticks (pale yellow). (C-D) Ribbon diagrams displaying B-factors of the structures of (C) the apo-PD-1N74G T76P A132V and (D) the apo-PD-1T76P A132V. (E-F) 2Fobs-Fcalc simulated-annealing composite-omit (5% atoms) electron density map contoured at 1.0 of (E) the apo-PD-1N74G T76P A132V and (F) the apo-PD-1T76P A132V structures. (G) Close-up view of the C D loop overlays of human apo-PD-1N74G T76P A132V (pale yellow) and pembrolizumab-bound PD-1 (PDB: 5GGS, chain Y, smudge green). Human PD-1 does not adopt an ordered μ-strand C”, and C and D are connected by an extended C D loop. (H) Ribbon diagram of human apo-PD-1N74G T76P A132V showing two neighboring PD-1 protomers (black and grey) in the crystal lattice. Pockets (pale green) and the FG loops (light blue) pack against each other between two protomers and form substantial crystal contacts.

**Figure S6.**
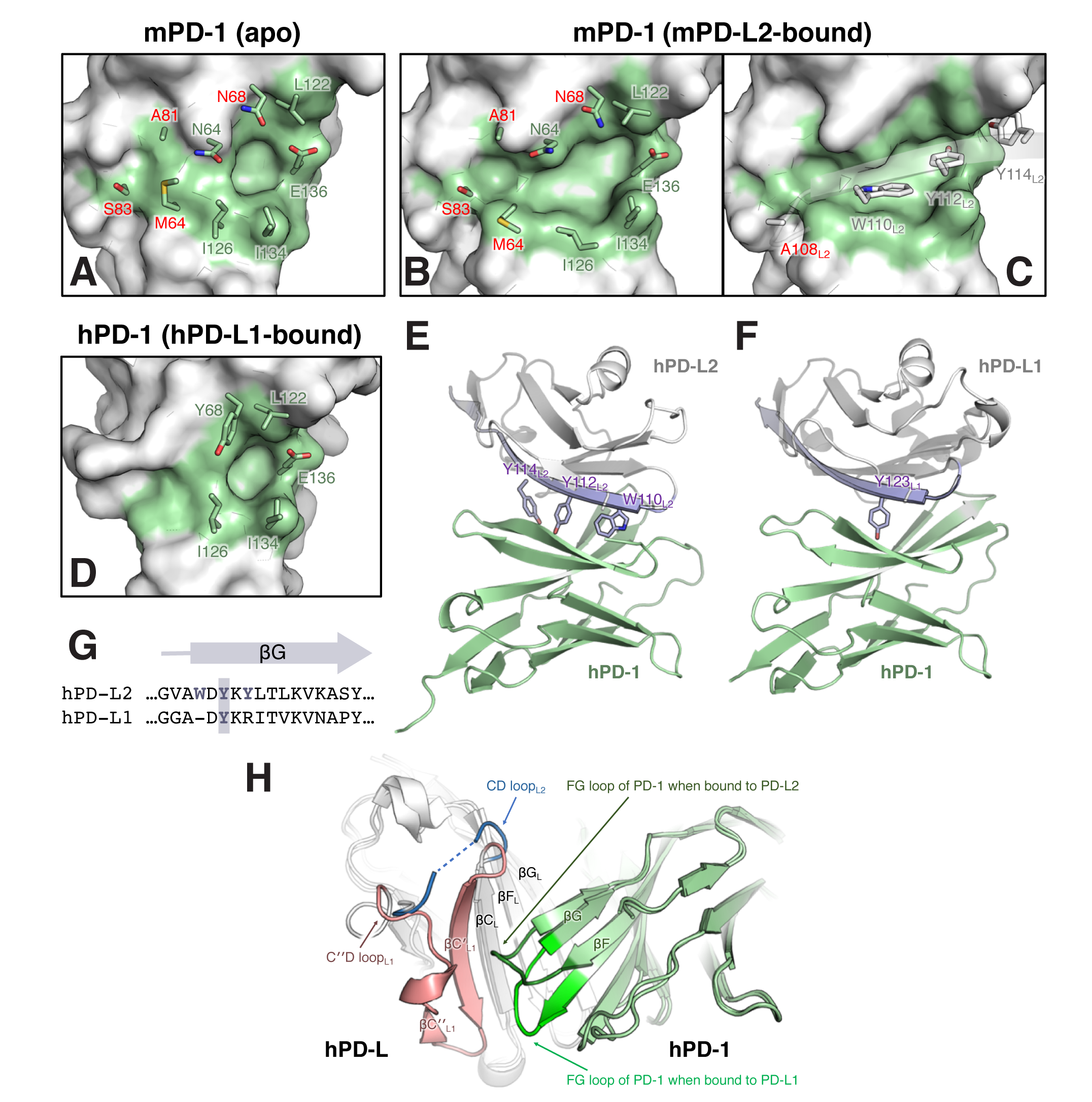
Pocket formation in murine PD-L2-bound murine PD-1 and human PD-L1-bound human PD-1, related to Figure 5. (A-D) Close-up views of space-filling models of (A) apo-murine PD-1 (PDB: 1NPU), and (B-C) murine PD-L2-bound murine PD-1 (PDB: 3BP5). In (C) a ribbon diagram of the G of PD-L2 is shown with PD-L2-interacting residues overlaid as sticks and labeled with an L2 subscript. Residues that are common (pale green) or different (red) between human and murine PD-1 are labeled. (D) human PD-L1-bound PD-1 (PDB: 4ZQK) overlaid with pocket residues shown as sticks. Pocket volumes are 220 Å3 for the murine pocket in the PD-1/PD-L2 structure and 80 Å3 for the human pocket in the PD-1/PD-L1 structure. (E) Ribbon diagram of the human PD-1/PD-L2 complex, with the sidechains of W110, Y112, and Y114 of PD-L2 shown as sticks and labeled with an L2 subscript. The sidechains of W110 and Y112 occupy the large pocket of PD-1 (Figure 5C). PD-1 (pale green), PD-L2 (grey), and G of the PD-L2 IgV domain (blue white) are highlighted. (F) Ribbon diagram of the human PD-1/PD-L1 complex (PDB: 4ZQK), with the sidechain of Y123 of PD-L1 shown as sticks and labeled with an L1 subscript. The sidechain of Y123 occupies the small pocket of PD-1 (Figure S6D). PD-1 (pale green), PD-L1 (grey), and G of the PD-L1 IgV domain (blue white) are highlighted. (G) Clustal Omega sequence alignment of G of the IgV domain between human PD-L2 and human PD-L1. G of PD-L1 IgV starts from residue D122. But G of PD-L2 IgV starts from residue W110, which is one residue earlier than the corresponding PD-L1 residue. Shaded tyrosines (Y112 of PD-L2 and Y123 of PD-L1) indicated a conserved aromatic residue. (H) Overlay of ribbon diagrams of the human PD-1/PD-L1 complex (PDB: 4ZQK) and the human PD-1/PD-L2 complex. PD-1 is colored in pale green, and PD-L1 and PD-L2 in grey. The IgV domain of PD-L1 adopts a C strand and a short C strand (pale red). The C strand form contacts with the FG loop of PD-1 (bright green). In contrast, the μ-strands C’ and C” are absence in the IgV domain of PD-L2. C is connected with the CD loop (blue). Part of the CD loop is flexible and unresolved in the crystal structure and a dash line is used to show connectivity. The FG loop of PD-1 (dark green) has a different conformation when bound to PD-L2. Key structural elements are labeled with an L1 subscript for PD-L1, an L2 subscript for PD-L2 or with an L subscript for both PD-L1 and PD-L2.

**Figure S7.**
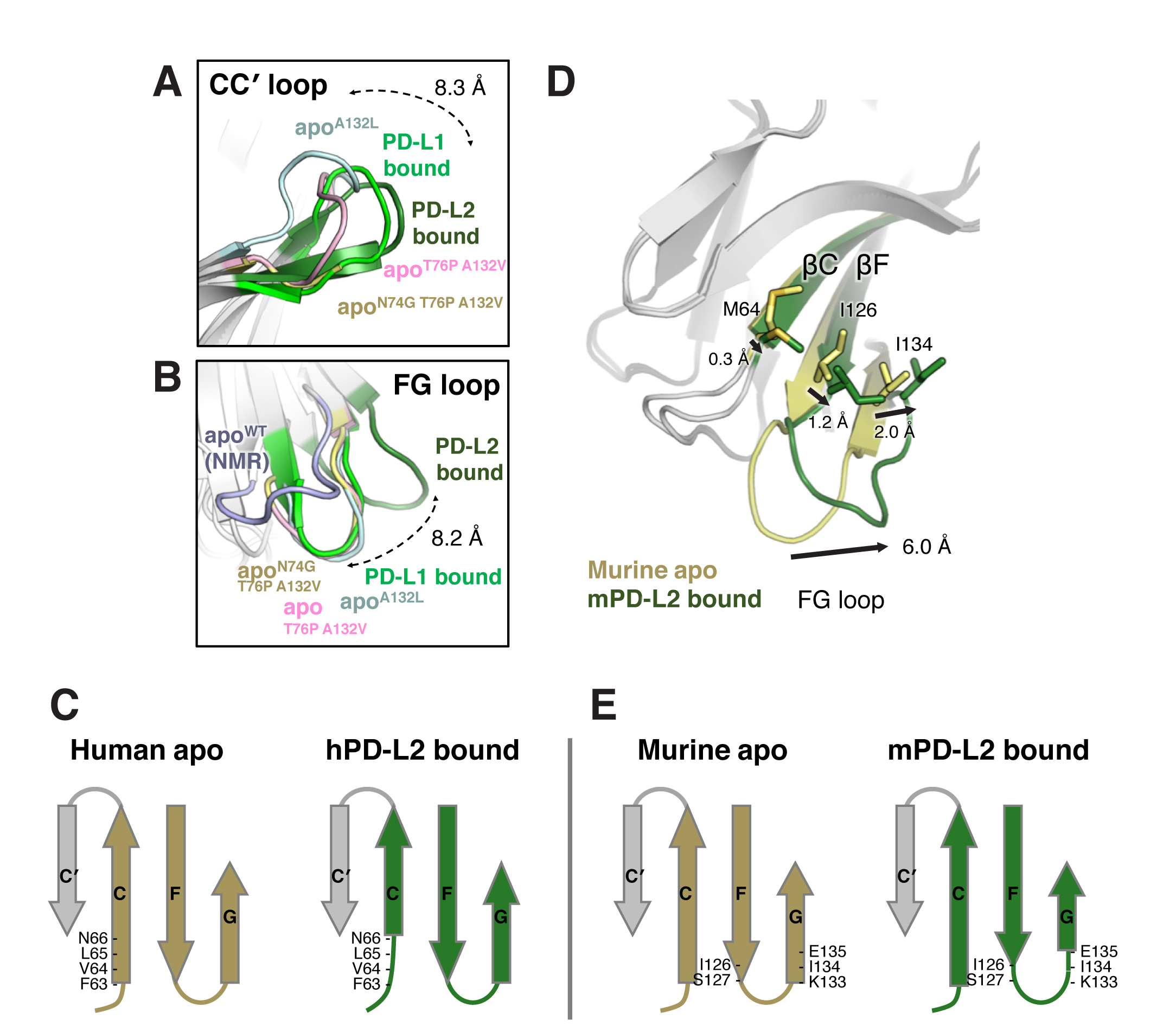
Systematic rearrangements in the PD-1 ligand-binding interface upon PD-L2 binding, related to Figure 6. (A-B) Overlays of ribbon diagrams of (A) the CC loops and (B) the FG loops from human PD-1 of apo-PD-1A132L (PDB: 3RRQ, cyan), apo-PD- 1T76P A132V (light pink), apo-PD-1N74G T76P A132V (pale yellow), PD-L1-bound PD-1 (PDB: 4ZQK, bright green), and PD-L2-bound PD-1N74G T76P A132V (dark green). (A) Dashed arrow depicts a C shift for Q75 of 8.3 Å from apo-PD-1 to PD-L2-bound PD-1N74G T76P A132V. In (B), an apo-human PD-1WT NMR structure (PDB: 2M2D, state 1, blue white) is also overlaid. Dashed arrow shows a C shift for P130 of 8.2 Å from apo-PD-1N74G T76P A132V to PD-L2-bound PD-1N74G T76P A132V. Residues 71-74 of the CC loop in apo-PD-1N74G T76P A132V are unresolved. Modest electron density for the mainchain atoms of these unresolved residues is observed in apo-PD-1T76P A132V. WT, wildtype; NMR, nuclear magnetic resonance. (C) Topology diagrams of the ligand-binding interface of human apo-PD-1N74G T76P A132V (left) and PD-L2-bound human PD-1N74G T76P A132V (right). C residues F63, V64, and L65 unfold into a coil upon PD-L2 binding. (D) Overlay of ribbon diagrams of murine PD-1 between apo-PD-1 (pale yellow) and PD-L2-bound PD-1 (PDB: 3BP5, dark green). A subset of pocket residues that undergo main-chain rearrangements are indicated with sticks. The FG loop shift of 6.0 Å in murine PD-1 was measured using the C of P130. (E) Topology diagrams of the ligand-binding interface of murine apo-PD-1 (left) and PD-L2-bound murine PD-1 (right). F residues I126 and S127, as well as G residues K133, I134, and E135, unfold into a coil upon PD-L2 binding.

**Figure S8.**
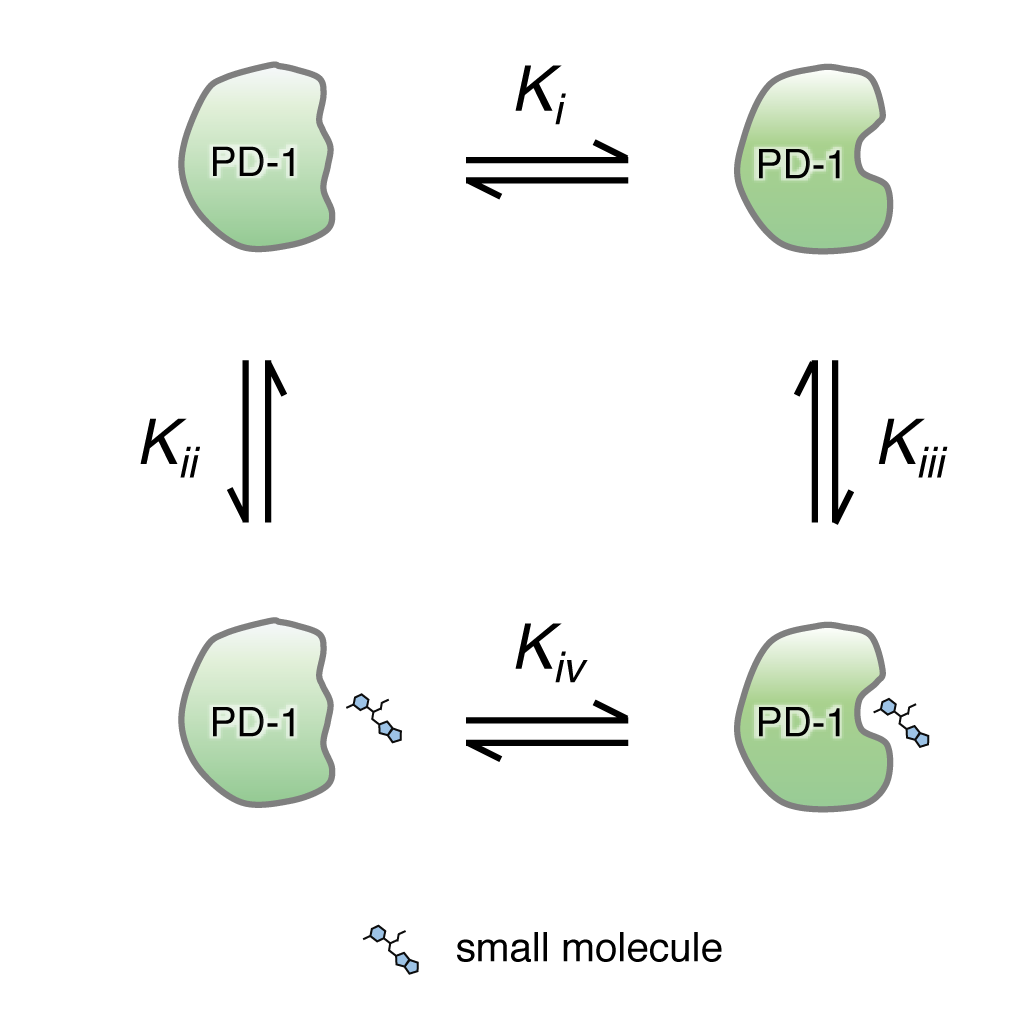
Thermodynamic cycle for binding of a small molecule to the PD-1 pocket. For clarity, only two of the states in the conformational ensemble of apo-PD-1 (top) are depicted. In apo-PD-1 the predominant species has a flat binding surface (Ki < 1). A pocket-binding drug will stabilize the PD-1 conformation containing the pocket (Kiii). Drug binding via an induced-fit mechanism (Kiv > 1) can also occur.

**Table S1.** Plasmids and yeast strains used in this study

| ***Saccharomyces cerevisiae* strain used for yeast surface display** |  |  |
| --- | --- | --- |
| **Strain** | **Genotype** | **Reference** |
| EBY100 | *MATa AGA1::P<sub>GAL1</sub>-AGA1::URA3 ura3-52 trp1 leu2Δ200 his3Δ200 pep4Δ::HIS3 prb1Δ1.6R can1 GAL* | (6) |
| **Plasmids for *Saccharomyces cerevisiae* expression** |  |  |
| **Plasmid** | **Description** | **Reference** |
| pST892 | pRS414 *P<sub>GAL1</sub>-AGA2-PD-1*(P21-E150) | This study |
| pST992 | pRS414 *P<sub>GAL1</sub>-AGA2-PD-1*(P21-E150) N74G | This study |
| pST993 | pRS414 *P<sub>GAL1</sub>-AGA2-PD-1*(P21-E150) T76P | This study |
| pST995 | pRS414 *P<sub>GAL1</sub>-AGA2-PD-1*(P21-E150) A132V | This study |
| pST1013 | pRS414 *P<sub>GAL1</sub>-AGA2-PD-1*(P21-E150) N74G T76P A132V | This study |
| **Plasmids for *Escherichia coli* protein expression** |  |  |
| **Plasmid** | **Description** | **Reference** |
| pST1132 | pET23d *PD-1*(N33-E150)-StrepII C93S N74G T76P A132V | This study |
| pST1167 | pET23d *PD-1*(N33-E150)-StrepII C93S T76P A132V | This study |
| **Plasmids for human Expi293F cell line protein expression** |  |  |
| **Plasmid** | **Description** | **Reference** |
| pST971 | pADD2 *PD-L1-Fc* | This study |
| pST972 | pADD2 *PD-L2-Fc* | This study |
| pST980 | pADD2 *Fc* | This study |
| pST981 | pADD2 *PD-1-Fc* | This study |
| pST982 | pADD2 *PD-1-Fc* N74G | This study |
| pST983 | pADD2 *PD-1-Fc* T76P | This study |
| pST985 | pADD2 *PD-1-Fc* A132V | This study |
| pST1008 | pADD2 *PD-1-Fc* N74G A132V | This study |
| pST1009 | pADD2 *PD-1-Fc* T76P A132V | This study |
| pST1010 | pADD2 *PD-1-Fc* N74G T76P A132V | This study |
| pST739 | pADD2 *PD-L1-His<sub>6</sub>* | This study |
| pST700 | pADD2 *PD-L2-His<sub>6</sub>* | This study |
| pST963 | pADD2 *PD-1-Fc* C93S | This study |
| pST964 | pADD2 *PD-1-Fc* C93S S71G P72G S73G N74G Q75G T76G D77G (CC' loop-mutant) | This study |
| pST965 | pADD2 *PD-1-Fc* C93S L128G A129G P130G K131G A132G Q133G (FG loop-mutant) | This study |
| pST966 | pADD2 *PD-1-Fc* C93S Y68A I126A I134A E136A (Pocket-mutant) | This study |
| pST1195 | pADD2 *PD-1*(N33-E150)-Ctag C93S N74G T76P A132V<br>N49S N58S N116D | This study |
| pST1228 | pADD2 *PD-1*(N33-E150)-Ctag N49D N58D N74D N116D | This study |
| pST1207 | pADD2 *PD-L2*(M1-Y123) N37D N64D | This study |
| pST1249 | pADD2 *PD-1-Fc* V64E | This study |
| pST1250 | pADD2 *PD-1-Fc* N66A | This study |
| pST1251 | pADD2 *PD-1-Fc* Y68A | This study |
| pST1252 | pADD2 *PD-1-Fc* Q75A | This study |
| pST1253 | pADD2 *PD-1-Fc* I126D | This study |
| pST1254 | pADD2 *PD-1-Fc* I134D | This study |
| pST1255 | pADD2 *PD-1-Fc* E136A | This study |
| pST1262 | pADD2 *PD-L2-His<sub>6</sub>* I103D | This study |
| pST1263 | pADD2 *PD-L2-His<sub>6</sub>* I105D | This study |
| pST1266 | pADD2 *PD-L2-His<sub>6</sub>* Y112A | This study |
| pST1267 | pADD2 *PD-L2-His<sub>6</sub>* Y114A | This study |

**Table S2.**
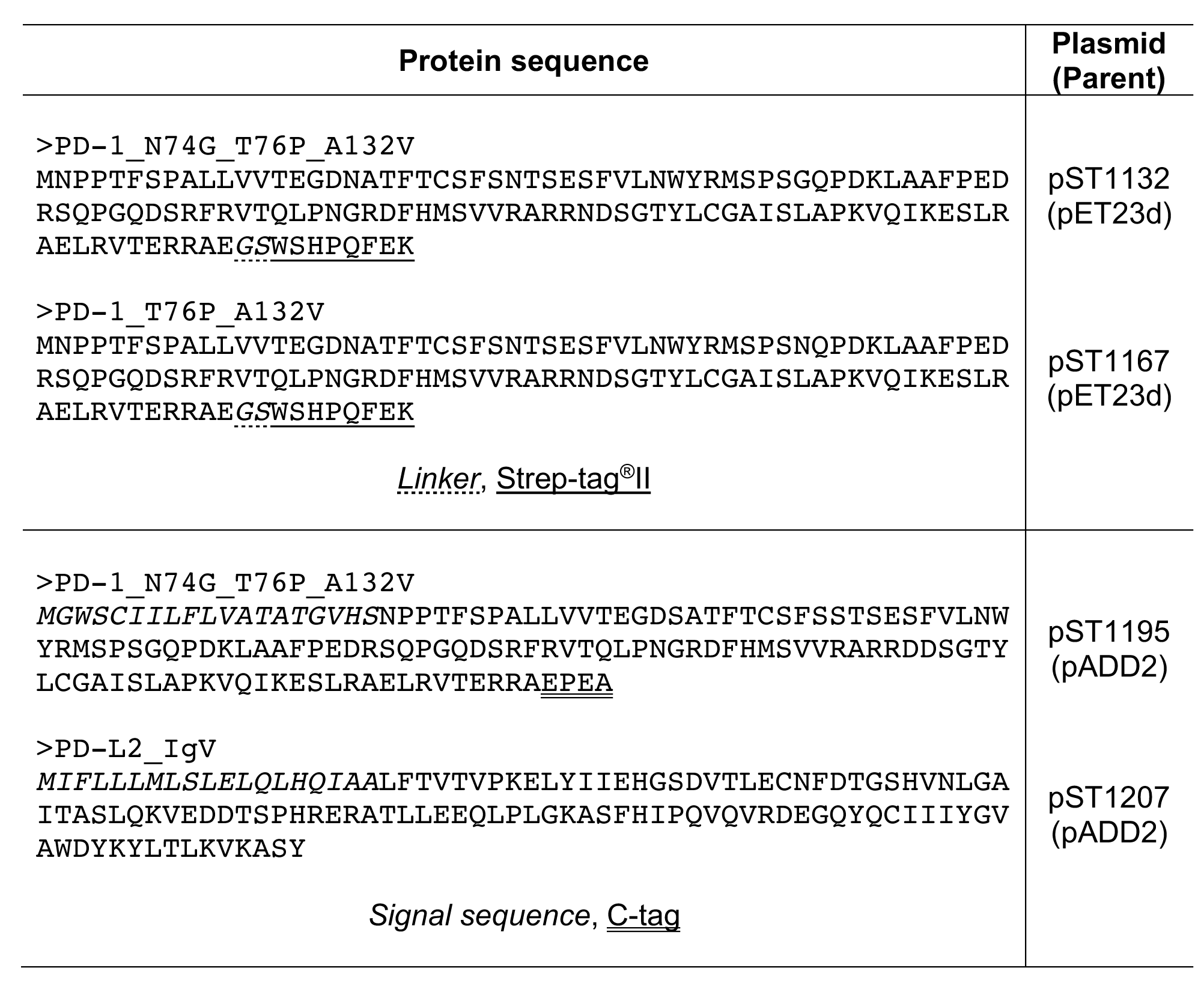
Protein sequences used for X-ray crystallography in this study

